## Supplementary Materials for "Defacing biases visual quality assessments of structural MRI"

#### General Supplementary Materials

***S1 Table. The IXI dataset was acquired at three different hospitals in London (UK).*** *Scanning parameters vary between sites. HH: Hammersmith Hospital, GH: Guy's Hospital and IoPPN: Institute of Psychiatry, Psychology & Neuroscience.*

| | HH | GH | IoPPN |
|---|---|---|---|
| Sample size | 185 | 322 | 74 |
| Field strength | 3T | 1.5T | 1.5T |
| Vendor & model (if available) | Philips Medical Systems Intera | Philips Medical Systems GyroScan Intera | General Electric |
| Repetition Time | 9.600 | 9.813 | |
| Echo Time | 4.603 | 4.603 | |
| Number of Phase Encoding Steps | 208 | 192 | |
| Echo Train Length | 208 | 0 | Details of scan parameter not available |
| Reconstruction Diameter | 240.0 | 240.0 | |
| Acquisition Matrix | 208 x 208 | - | |
| Flip Angle | 8.0 | 8.0 | |

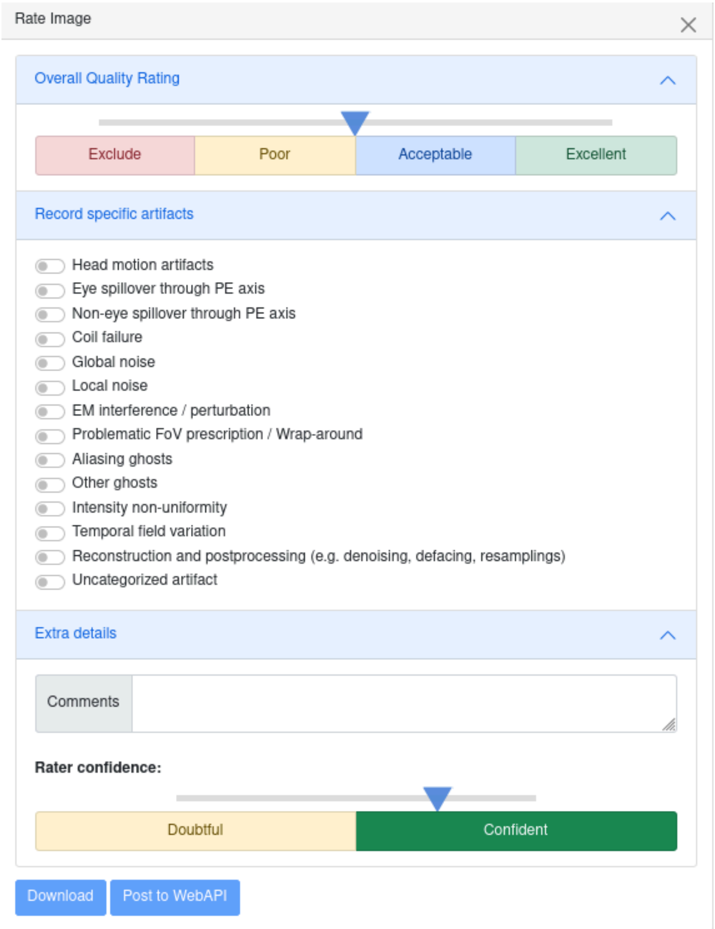

***S1 Figure. The updated MRIQC rating widget to assign quality grades using a slider.*** *The updated widget ranges from 1 to 4 and produces interval ratings. Categories are indicated as hints, but the actual ratings are fine-grained (with interval steps of 0.05). Additionally, we added a field to insert comments and a slider to indicate the rater's confidence on a scale from 0 to 1 with two referencing categories below. The raters' confidence and the selected artifacts lists were recorded and shared for future exploration (see Data and code availability statement). Still, they were not employed in any of the analyses presented in this report.*

### S1. Methodological clarifications

#### S1.1. Interpretation of BA plots, LoA, and CI of the bias

Bland & Altman indicated that even though a bias may not be significant, that doesn't mean that the bias is clinically irrelevant. For example, if the bias between two blood pressure devices is 1.5 mm Hg with a 95% CI of –0.2 to 3.2, since zero is within this interval, there is no significant bias. The 95% LoA might be –5.0 to 8.0 mm Hg, meaning most measurements differ by at most about 5 to 8 units. A clinician will judge if that range is acceptable. The reverse situation may also be true: a significant bias can be clinically irrelevant, which is what occurred to the vast majority of IQMs (Figure S16). Suppose the two devices showcase a bias of 3 mm Hg, with 95% CI of 2.999 to 3.001 mm Hg—i.e., significant bias,—and a 95% LoA of 2.95–3.05. The two devices are interchangeable despite the significant bias because they agree substantially—provided the 0.1 mm span of LoA is not clinically meaningful. Indeed, to use one or another, we would need to account for the 3±0.001 mm Hg systematic bias between the two devices.

#### S1.2. Effect size estimation of (M)ANOVA models

Because effect sizes of type partial eta-squared ($\eta^2$) are most commonly reported for ANOVAs, we converted the effect size of type f generated by the sensitivity analysis with *G*Power* (Faul et al. 2009) to type partial $\eta^2$ using:

$$\eta_p^2 = \frac{f^2}{1+f^2} \quad \text{(Equation S1).}$$

In our pilot study, we computed the effect size associated with the bias on human quality assessment as indicated in Equation S2. Note that we pooled the ratings from both raters before applying this formula.

$$d = \frac{|mean(\,nondefaced\,ratings) - mean(defaced\,ratings)|}{std(all\ ratings)} = \frac{0.45}{0.72} = 0.62 \quad \text{(Equation S2)}$$

We had to convert Cohen's d, reported in previous studies, to Cohen's f because the ANOVA family of tests we used in this pre-registration reported Cohen's f. When comparing two groups with equal sample sizes, the conversion between Cohen's d and Cohen's f is simply:

$$f = \frac{d}{2} \quad \text{(Equation S3)}$$

To compute the effect size associated with the MANOVA test performed on the IQMs of the pilot study, we used Equations S4 and S5:

$$\eta_p^2 = \frac{numDF * F}{numDF * F + denDF} = \frac{5*0.0721}{5*0.0721 + 14} = 0.025 \quad \text{(Equation S4)}$$

$$f = \sqrt{\left(\frac{\eta_p^2}{1-\eta_p^2}\right)} = 0.16 \quad \text{(Equation S5)}$$

### S2. Exploratory analysis of the manual ratings

#### S2.1. Understanding the outcomes of manual assessment

***S2 Table. Defacing-derived bias was larger and more consistent than intrinsic intra-rater variability.*** *To compare intra-rater reliability to the reliability with respect to the defacing condition, we summarized in this table the reliability measures stemming from the comparison of ratings in nondefaced/defaced image pairs (BA plot on Fig. S2) as well as the comparison of ratings between the first and second presentation of repeated images (BA plot on Fig. S7). Note that we shuffled the presentation order of defaced and nondefaced images, which enabled us to distinguish biases introduced by defacing from those by the order of presentation. We quantified the reliability using two metrics: the bias computed as the mean difference between image pairs and using agreement two-way random effects intra-class correlation (ICC(2,1)) implemented with the irr R package (Gamer, Lemon, and Singh 2019; R Core Team 2021). Not all raters were equally biased by defacing, but the sign of the bias remained consistent, meaning that all raters typically overestimated the quality of defaced images. However, note that the defacing bias of Raters 2 and 4 were not significant as the bias 95% confidence interval (CI) contained zero. Similarly, not all raters were equally consistent with themselves when assessing the quality of the same image, but in this case, the sign of the bias varied. Only Rater 1 showed significant intra-rater variability as indicated by the bias 95% CI not containing zero. Regarding the ICC, all raters presented intra-rater reliability considered good (Cicchetti and Sparrow 1981) and higher than the inter-rater reliability (0.542; see section S4.2). The ICC related to defacing was consistently lower than the intra-rater ICC (except for Rater 4). As observed in Figs. 2 and S3, Rater 1—the most experienced—presented the largest bias and the lowest reliability.*

| | | Rater 1 | Rater 2 | Rater 3 | Rater 4 | All raters |
|---|---|---|---|---|---|---|
| $\mathbb{E}[\Delta_{ndef\text{-}def}]$ [95% CI] | Defacing | -0.12* [-0.14, -0.08] | -0.01 [-0.04, 0.02] | -0.06* [-0.08, -0.03] | -0.02 [-0.03, 0.004] | -0.05* [-0.06, -0.03] |
| | Intra-rater | 0.11* [0.017, 0.21] | -0.009 [-0.09, 0.08] | -0.07 [-0.15, 0.0005] | 0.05 [-0.12, 0.11] | 0.02 [-0.02, 0.06] |
| ICC (2,1) [95% CI] | Defacing | 0.539 [0.434, 0.629] | 0.62 [0.532, 0.694] | 0.68 [0.603, 0.745] | 0.734 [0.667, 0.769] | 0.694 [0.658, 0.727] |
| | Intra-rater | 0.609 [0.449, 0.731] | 0.676 [0.536, 0.779] | 0.746 [0.63, 0.83] | 0.704 [0.574, 0.799] | 0.735 [0.68, 0.782] |

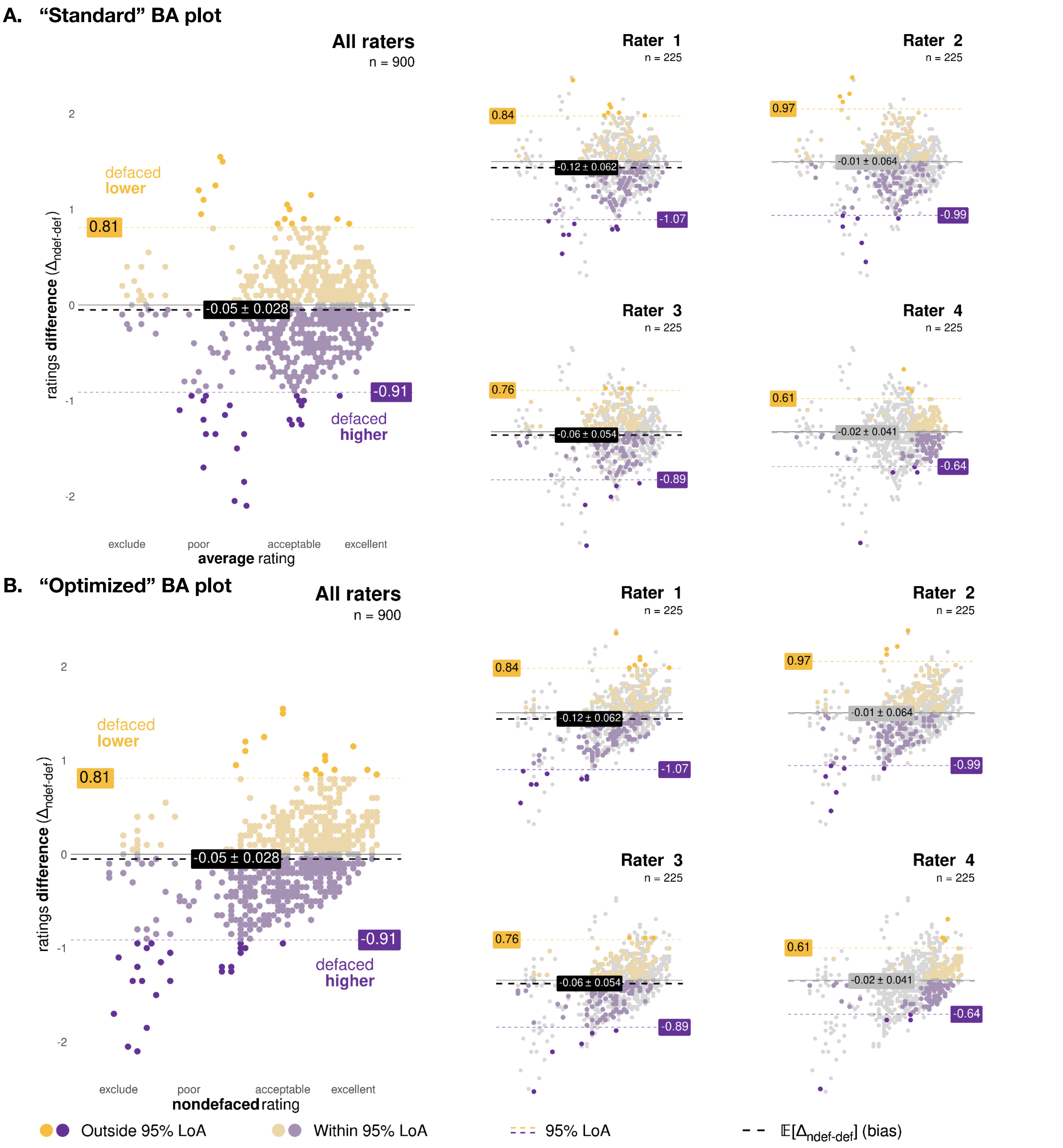

***S2 Figure. Optimizing BA plots to overcome the wrapping effect of our quality scale.*** *When we screened the "standard" BA plots, where the x-axis represents the mean of the two measurements, and the y-axis represents their difference (Panel A, top), we noticed an apparent "wrapping" effect of the manual ratings, revealing that the images with* $\Delta_{ndef\text{-}def}$ = [1.75, 2.5]*—corresponding to a "poor" through "acceptable" range of the MRIQC slider (c.f., Fig. S1),—presented the largest* $\Delta_{ndef\text{-}def}$ *differences. Our "optimized" BA plots (Panel B, bottom; and Fig. 2 of the main text) use the nondefaced rating on the x-axis rather than the mean. The optimization unveils how the bias direction shifts predictably based on the nondefaced rating. A high-rated nondefaced instance is unlikely to receive a higher rating in the defaced condition because the ratings are bounded in the [1,4] interval. Correspondingly, a low-rated nondefaced instance is unlikely to receive a lower rating when defaced.*

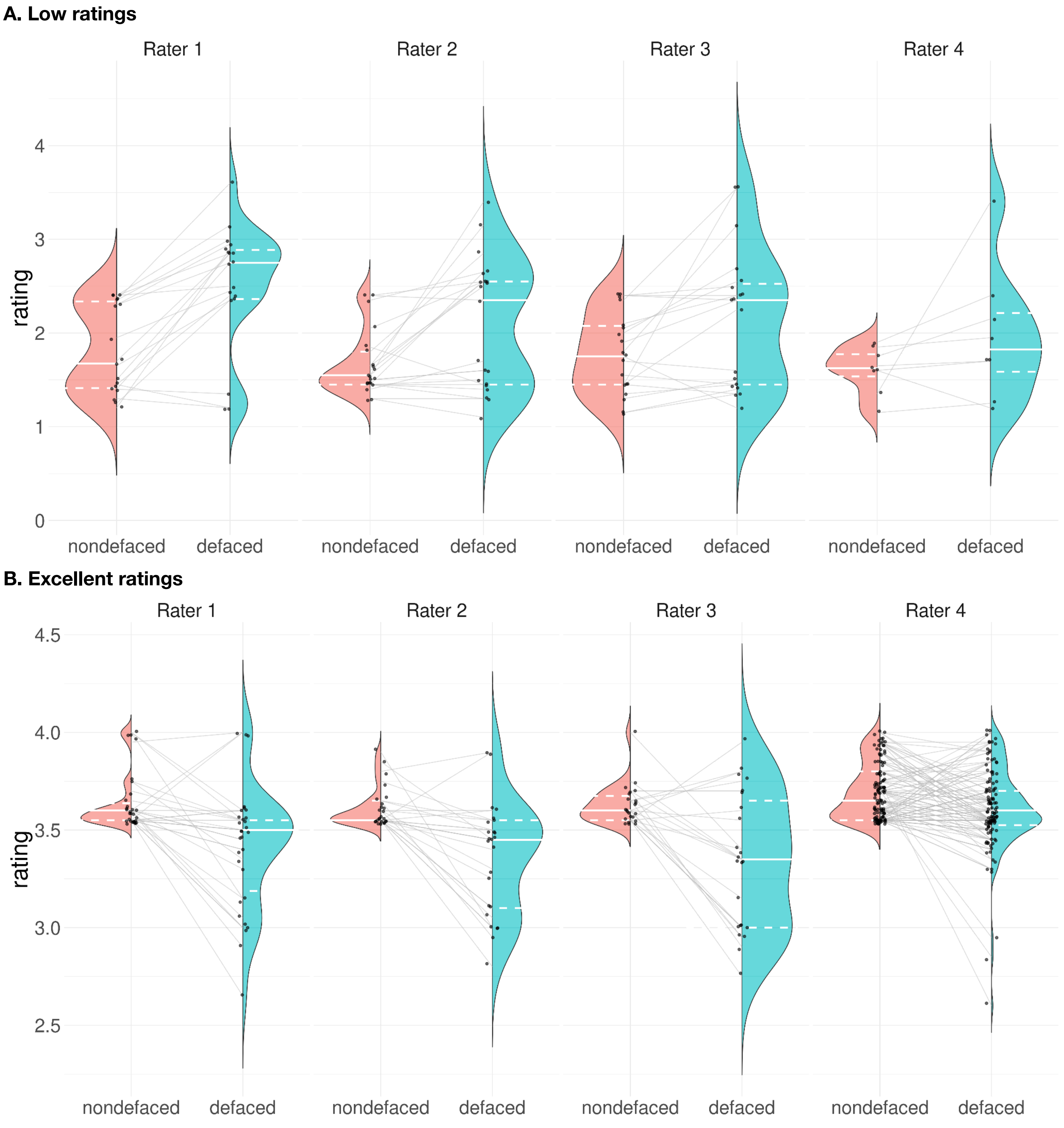

***S3 Figure. Exploring the variability introduced at the tails of the nondefaced ratings distribution.*** *We employed the "test-retest violin plot" of Fig. 3 to investigate the effect of defacing on low-rated nondefaced images (Panel A; rating below 2.45; 18, 21, 19, 8 nondefaced/defaced rating pairs per rater, respectively); and high-rated instances (Panel B; >3.55; 30, 25, 23, 115 rating pairs per rater, respectively). Panel A suggests that Rater 1 (the most experienced) was the most sensitive to the defacing operation with a more apparent pattern of nondefaced-defaced trajectories with positive slope between the half-violins, with the gap often reaching 1 unit (equivalent to switching category in the quality appreciation scale). Excellent ratings show overall negative trajectories, but the range of variation is limited to 0.5 units, with a few exceptions that reach a 0.75 unit change.*

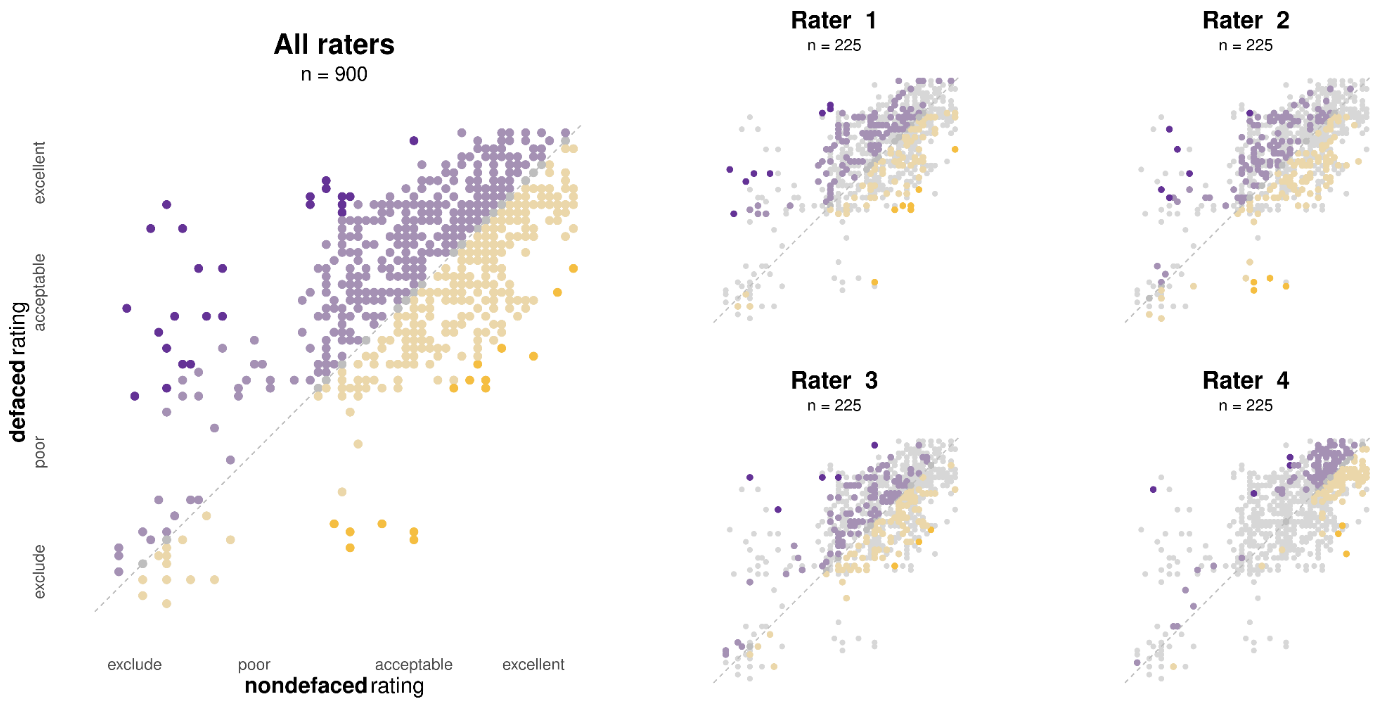

***S4 Figure. Exploring the correlation between nondefaced and defaced ratings.*** *Reliable ratings ($\Delta_{ndef\text{-}def} \approx 0$) are located near the dashed line of slope one when presenting nondefaced vs. defaced scatter plots. Instances above this line are presented in purple, reflecting that the defaced rating was higher than its corresponding counterpart ($\Delta_{ndef\text{-}def} < 0$). Conversely, instances below the equality line are represented in yellow ($\Delta_{ndef\text{-}def} > 0$). This color convention is consistent with Figs. 2 (main text) and S2. Further supporting Fig. 2 (main text), this figure highlights that if a nondefaced image was rated as low-quality, the assessment of the defaced image quality tended to be over-optimistic (represented with darker purple dots).*

#### S2.2. Inter-rater reliability

The inter-rater reliability was 0.542 (95% CI = [0.497, 0.587]), which corresponds to a moderate agreement between raters (Cicchetti and Sparrow 1981). We calculated this reliability using consistency two-way random effects intra-class correlation (ICC(2,1)) implemented with the irr R package (Gamer, Lemon, and Singh 2019; R Core Team 2021).

#### S2.3. Intra-rater reliability

In both conditions, 40 subjects were presented to the raters twice to assess the intra-rater reliability. As the subject identifiers were obfuscated, the raters were unaware that they rated images twice. The order of presentation of all images was random. We generated several plots to visualize the intra-rater variability and quantified it using the intraclass correlation coefficient (ICC).

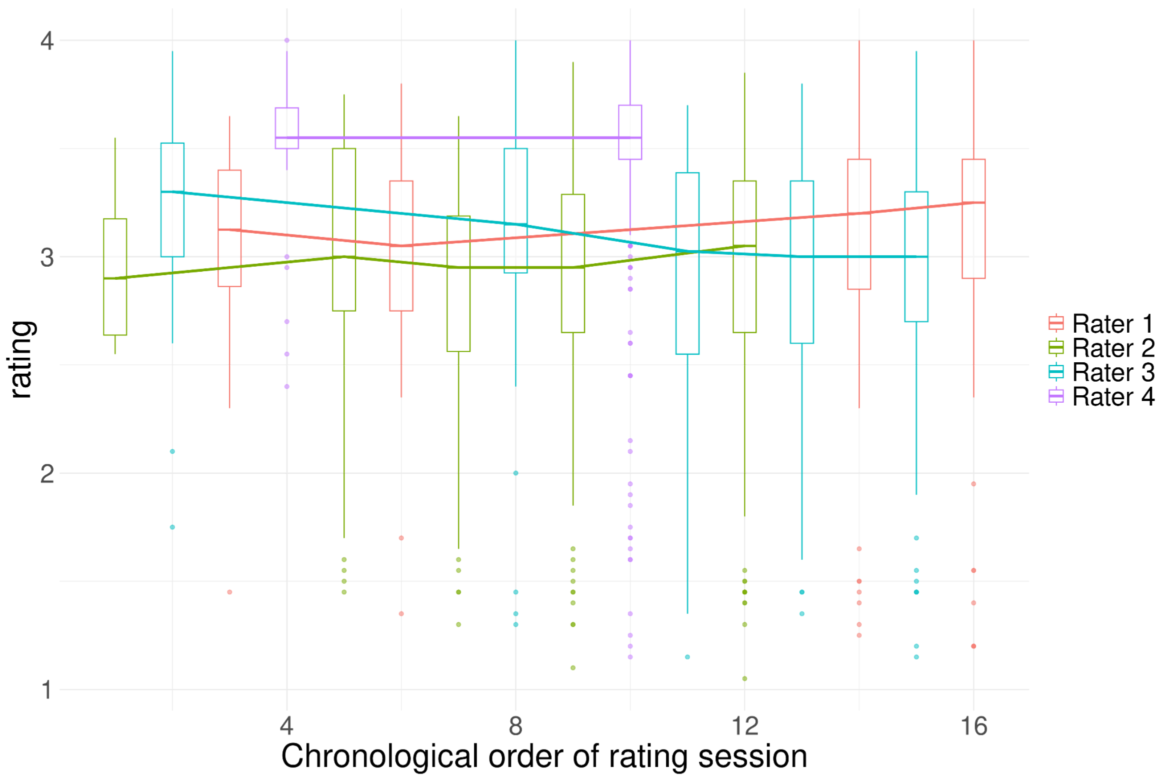

***S5 Figure. Ratings' evolution throughout different rating sessions.*** *We identified groups of ratings that were assigned during the same rating session by the same rater, and for each session, we plotted the rating distribution in a boxplot. Identifying the rating session was possible because our data frame is organized chronologically, meaning the ratings are listed in the order they were assigned. Rater 4 assessed most of the 450 images in one session, which may explain why they tended to give similar ratings to all images (shown by the narrow standard deviation of his rating distribution). Moreover, Rater 3 became more critical of image quality over time. The central dash in the boxplot corresponds to the median, the box limits correspond to the 25$^{th}$ and 75$^{th}$ percentiles, and the whiskers extend to the largest, respectively smallest, value no further than 1.5 times the inter-quartile range.*

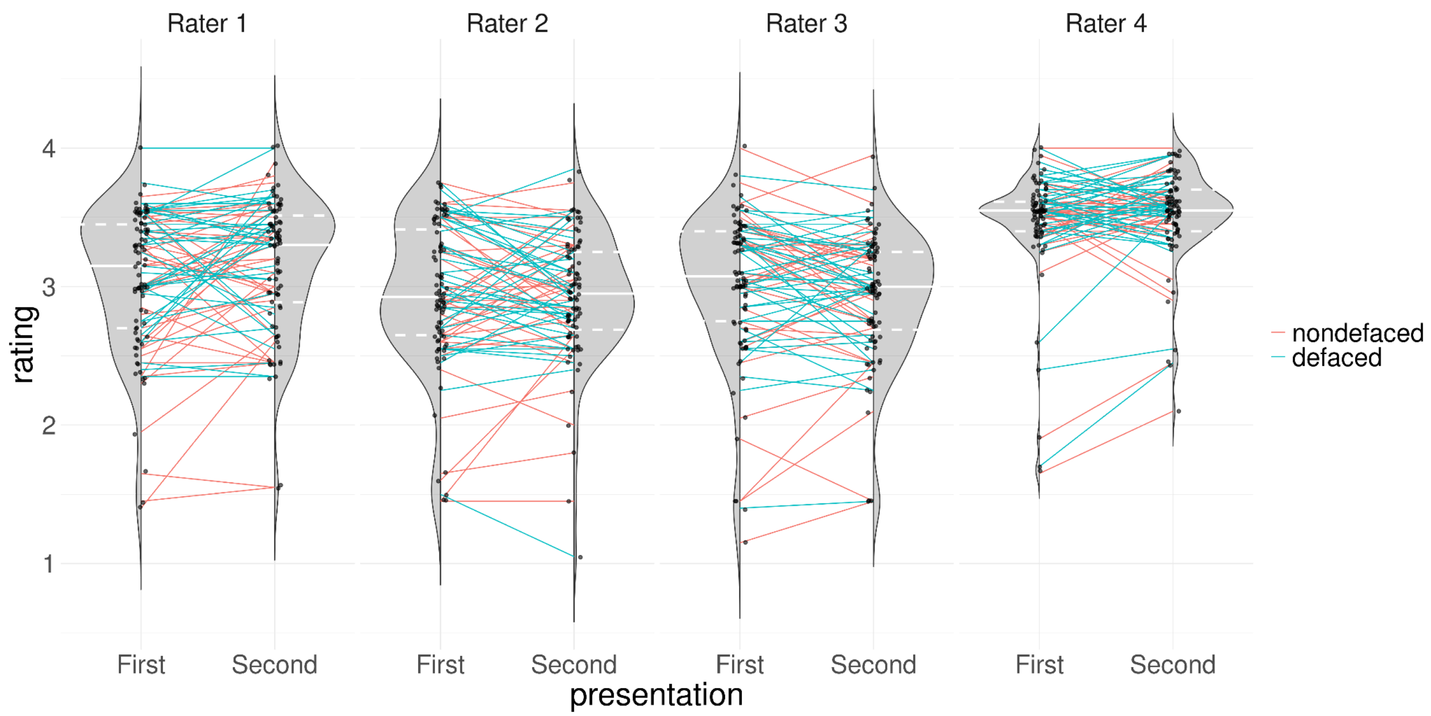

***S6 Figure. Test-retest violin plots comparing the rating assigned to the first and second presentation of the repeated images.*** *Mapping lines link the two ratings of a single instance and are color-coded, indicating whether the image was defaced (blue) or nondefaced (red). The distribution fit displays the median with a solid-white line, while dashed lines represent the 25% and 75% quantiles. We did not identify consistent patterns associated with the defacing condition, as suggested by all first and second distributions being centered at similar locations. Rater 1 gave the second image presentation a better rating, as visible in the median difference between the two violins. Rater 3 typically gave lower ratings for the second image presentation.*

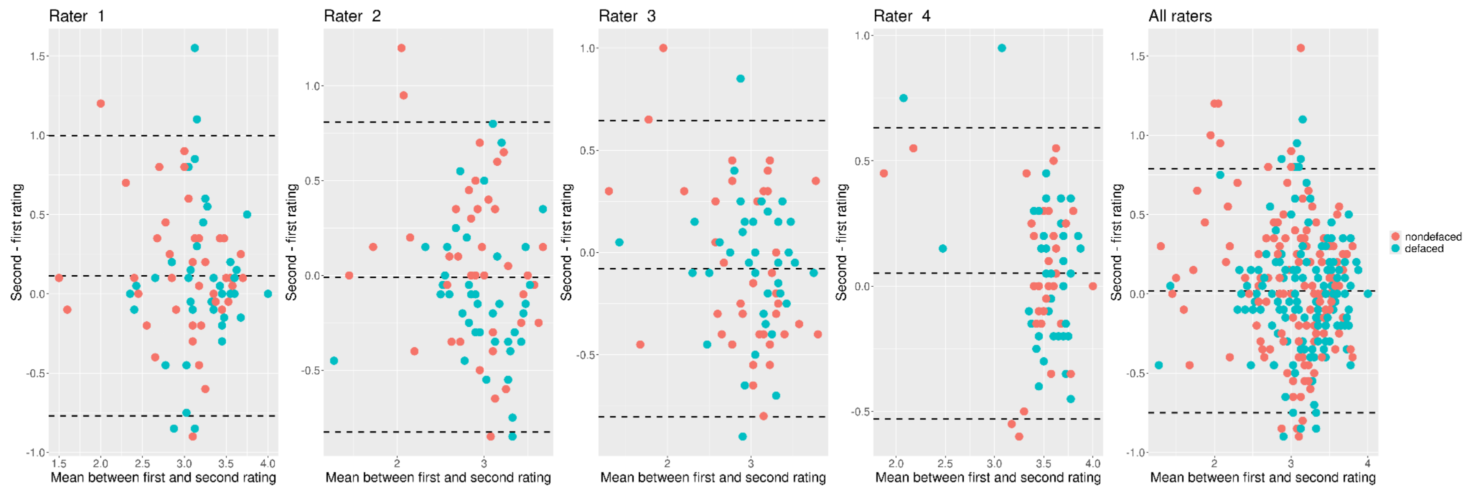

***S7 Figure. The test-retest exploration did not reveal consistent intra-rater biases.*** *Raters 1 and 4 showcased positive mean differences, which may be due to raters becoming decreasingly sensitive to the artifacts in the dataset over time, e.g., due to fatigue. On the contrary, Rater 3 seemed to have become better at spotting issues over time, as they presented a negative mean difference. Rater 2 presented a bias close to zero, meaning they gave a consistent rating between the first and second image presentations. The bias, displayed with a solid black line, corresponds to the mean difference in rating between the first and second presentation of the same image. The values of the biases are reported in Table S2. Dashed lines correspond to the 95% limits of agreement computed as bias ±1.96·SD, with SD being the standard deviation of the differences between test and retest.*

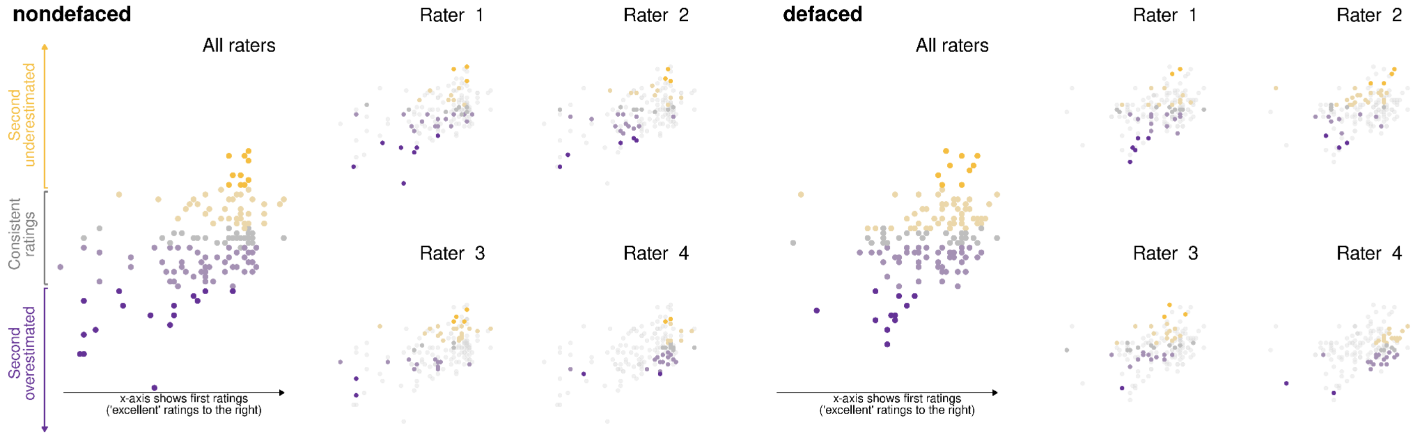

***S8 Figure. Optimized BA plots of intra-rater variability by defacing condition.*** *The x-axis corresponds to the first rating assigned, while the y-axis corresponds to the difference between the second and first rating. Only the images that were repeated twice were considered for this plot. The color code is the same as in Fig. 2 of the main manuscript. The constraint that our rating scale imposes on the quality score is still visible; the high-rated images were assigned a similar or lower grade in the second presentation, whereas low-rated images were assigned a similar or higher grade in the second presentation. The defaced distribution is also compressed towards the right instead of the nondefaced distribution, which spreads further towards low ratings. This observation agrees with our conclusion that lower ratings received substantially better after defacing from Fig. 2.*

#### S3. Analysis of the manual ratings

##### S3.1. Repeated-measures ANOVA

We checked whether the data were normally distributed within each subgroup, which was defined by the intersection between rater and defacing conditions by two means. First, we ran Shapiro-Wilk normality tests (Shapiro and Wilk 1965) and reported the results in Table S3. Secondly, we visualized Q-Q plots (Wilk and Gnanadesikan 1968) presented in Fig. S9. We also checked the normality of each subgroup using only the poor-quality and excluded images. As pre-registered, we also assessed that the sphericity assumption was not met in the full-dataset and low-ratings tests (Table S4). However, the violation of the sphericity assumption disappeared when Rater 4 was excluded, providing further evidence that the rating distribution of this rater was not comparable to the other three. Acknowledging that violating the normality and sphericity assumptions would decrease the sensitivity of the test, we ran the rm-ANOVA on the three sets of data (full dataset, only Raters 1, 2, and 3, and only lower-quality scans) to complement the LMEs analysis We found defacing a significant effect after correction for multiple comparisons in the full-sample and the Rater 4-excluded tests (see Table S4 for full reporting and interpretation).

***S3 Table. Normality tests.*** *The Shapiro-Wilk normality test (Shapiro and Wilk 1965) indicated that ratings were not normally distributed, except for the lower ratings of Rater 3 (nondefaced ratings) and Rater 4 (both conditions).*

| | | Rater 1 | | Rater 2 | | Rater 3 | | Rater 4 | |
|---|---|---|---|---|---|---|---|---|---|
| | | W | p | W | p | W | p | W | p |
| All ratings | nondefaced | 0.896 | $4{\cdot}10^{-10}$ | 0.910 | $3{\cdot}10^{-9}$ | 0.908 | $2{\cdot}10^{-9}$ | 0.649 | $<2{\cdot}10^{-16}$ |
| | defaced | 0.902 | $10^{-9}$ | 0.893 | $6{\cdot}10^{-10}$ | 0.914 | $6{\cdot}10^{-9}$ | 0.649 | $<2{\cdot}10^{-16}$ |
| Low ratings only (<2.45) | nondefaced | $4{\cdot}10^{-10}$ | $4{\cdot}10^{-4}$ | 0.833 | 0.001 | 0.912 | 0.06 | 0.929 | 0.509 |
| | defaced | 0.604 | $7{\cdot}10^{-6}$ | 0.833 | 0.001 | 0.745 | $5{\cdot}10^{-5}$ | 0.925 | 0.475 |

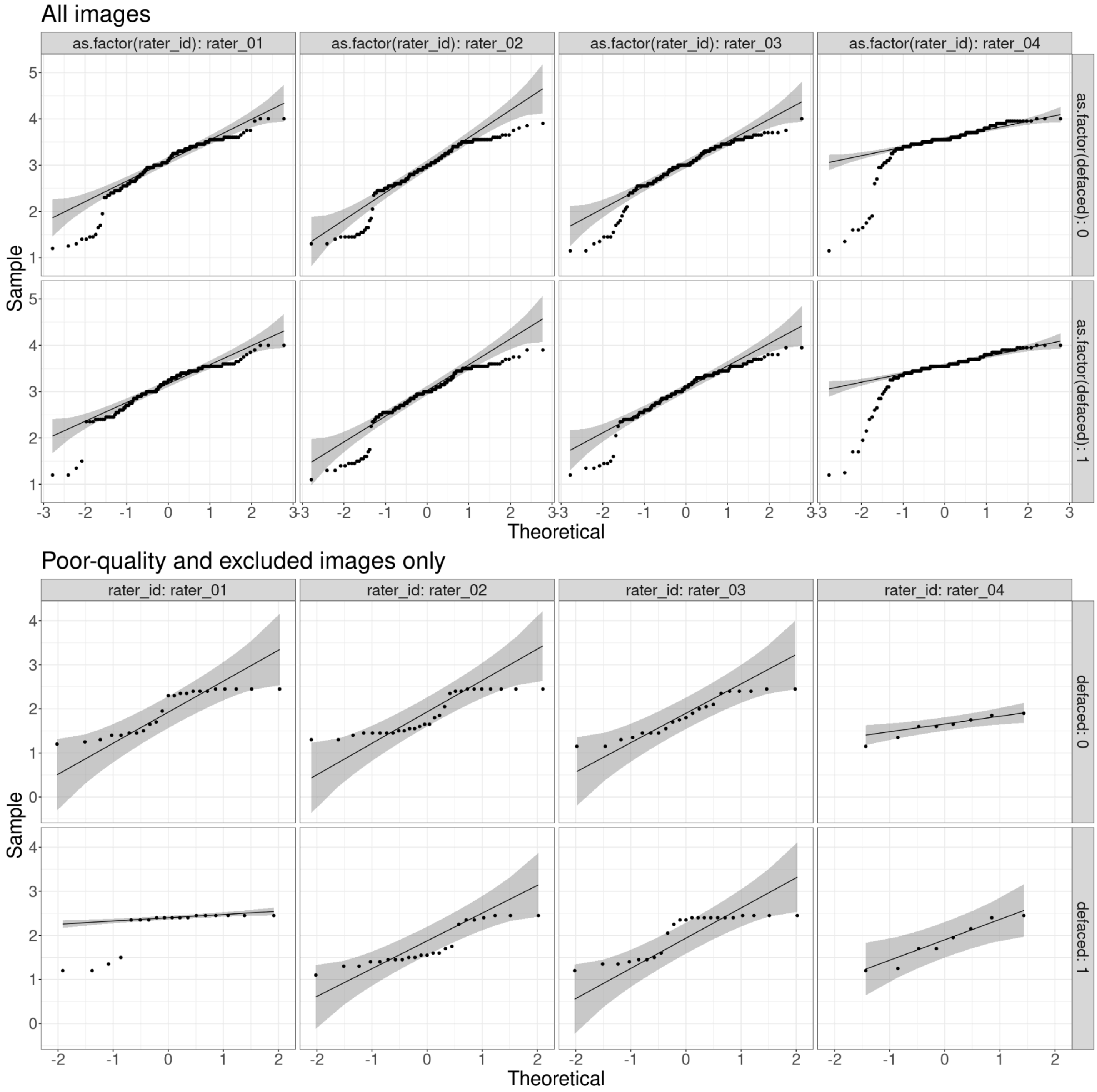

***S9 Figure. Check for rating normality using a Q-Q plot.*** *The Q-Q plot is a graphical method to compare probability distribution by plotting their quantiles against each other (Wilk and Gnanadesikan 1968). The reference line represents perfect normality. When some ratings fall outside the shaded region, the data are not normally distributed. When considering all images, none of the subgroups was normally distributed. In contrast, low ratings deviate much less from normality, except for ratings assigned by Rater 1 in the defaced condition.*

***S4 Table. Results of repeated-measures ANOVA.*** *We ran three rm-ANOVA: one including all ratings, one excluding Rater 4, and one including only ratings below 2.45 corresponding to images rated as poor quality or excluded. $p_{corrected}$ correspond to p-values corrected for multiple comparisons controlling for false discovery rate (Benjamini and Hochberg 1995). The p-values related to the defaced effect were corrected, considering the p-values from the three likelihood ratio tests (c.f., Table S7). We report the rm-ANOVA effect sizes in the form of generalized eta-squared (ges) and partial eta-squared (pes; $\eta_p^2$). The effect size linked to the defacing factor of the rm-ANOVA with low ratings being larger than that of the rm-ANOVA with all raters (0.765 versus 0.037) confirmed that low ratings were the ones most influenced by the process of defacing. The non-significance of the rm-ANOVA with low ratings despite bigger effect size can be explained by the small sample available for this test (152 ratings).*

| | | **All raters (pre-registered)** | | | **Without Rater 4 (exploratory)** | | | **Low ratings only (exploratory)** | | |
|---|---|---|---|---|---|---|---|---|---|---|
| | | Defaced | Rater | Interaction | Defaced | Rater | Interaction | Defaced | Rater | Interaction |
| **ANOVA** | sample size | 1480 | | | 1110 | | | 152 | | |
| | DFn | 1 | 3 | 3 | 1 | 2 | 2 | 1 | 3 | 3 |
| | DFd | 184 | 552 | 552 | 184 | 368 | 368 | 3 | 9 | 9 |
| | F | 7.08 | 158.904 | 1.826 | 7.745 | 15.023 | 1.468 | 9.768 | 3.162 | 1.167 |
| | p | 0.008 | $3 \cdot 10^{-7}$ | 0.141 | 0.006 | $5 \cdot 10^{-7}$ | 0.0232 | 0.052 | 0.078 | 0.375 |
| | $p_{FDR}$ | 0.016 | $9 \cdot 10^{-74}$ | 0.348 | 0.016 | $8 \cdot 10^{-7}$ | 0.348 | 0.052 | 0.078 | 0.375 |
| | p<.02 | * | *** | | * | *** | | | | |
| | ges | 0.002 | 0.155 | 0.001 | 0.003 | 0.015 | 0.001 | 0.097 | 0.311 | 0.078 |
| | pes ($\eta_p^2$) | 0.037 | 0.463 | 0.010 | 0.04 | 0.075 | 0.008 | 0.765 | 0.513 | 0.280 |
| **Mauchly's Test for Sphericity** | W | | 0.942 | 0.901 | | 0.984 | 0.980 | | 0.1430 | $1 \cdot 10^{-4}$ |
| | p | | 0.054 | 0.002 | | 0.226 | 0.160 | | 0.699 | 0.022 |
| | p<.05 | | | * | | | | | | * |
| **Sphericity Corrections** | GGe | | 0.963 | 0.938 | | 0.984 | 0.981 | | 0.511 | 0.428 |
| | DF [GG] | | 2.89, 531.79 | 2.81, 517.66 | | 1.97, 362.16 | 1.96, 360.84 | | 1.53, 4.6 | 1.28, 3.85 |
| | p[GG] | | $1 \cdot 10^{-71}$ | 0.145 | | $6 \cdot 10^{-7}$ | 0.232 | | 0.139 | 0.366 |
| | p[GG]<.05 | | * | | | * | | | | |
| | HFe | | 0.980 | 0.954 | | 0.995 | 0.991 | | 0.937 | 0.610 |
| | DF[HF] | | 2.94, 541.18 | 2.86, 526.52 | | 1.99, 366.05 | 1.98, 364.69 | | 2.81, 8.44 | 1.83, 5.49 |
| | p[HF] | | $1 \cdot 10^{-73}$ | 0.144 | | $6 \cdot 10^{-71}$ | 0.232 | | 0.084 | 0.372 |
| | p[HF]<.05 | | * | | | * | | | | |

Interaction denotes the interaction between the defaced condition and the rater
sample size: number of ratings considered in the model

***S5 Table. Results of running one repeated-measures ANOVA for each rater separately.*** *$p_{FDR}$ corresponds to p-values for multiple comparisons controlling for false discovery rate (Benjamini and Hochberg 1995). No rater showed a significant bias due to defacing after correction for multiple comparisons. The effect sizes are reported in generalized eta-squared form (ges) and partial eta-squared form (pes $\eta_p^2$).*

| Rater ID | Sample size | Effect | DFn | DFd | F | p | $p_{FDR}$ | p<.02 | ges | pes ($\eta_p^2$) |
|---|---|---|---|---|---|---|---|---|---|---|
| Rater 1 | 370 | defaced | 1 | 184 | 7.655 | 0.006 | 0.024 | | 0.009 | 0.04 |
| Rater 2 | | | | | 0.179 | 0.673 | 0.816 | | $19 \cdot 10^{-5}$ | 0.000971 |
| Rater 3 | | | | | 3.87 | 0.051 | 0.102 | | 0.003 | 0.021 |
| Rater 4 | | | | | 0.054 | 0.816 | 0.816 | | $3 \cdot 10^{-5}$ | 0.000295 |

sample size: number of ratings considered in the model

#### S3.2. Linear mixed-effects models

***S6 Table. Model fit indices of LMEs with ("alternative") and without ("baseline") defacing as a fixed effect.*** *Both the baseline and alternative models included subject and rater as random effects. We fitted both models on three data subsamples: all ratings, only ratings of Raters 1, 2, and 3, and only low ratings (<2.45) corresponding to images rated as poor quality or excluded. In all three tests, the alternative model presented a lower AIC, lower BIC (except for the LME with all raters and with only poor-quality scans), higher log-likelihood, and lower deviance, indicating that including defacing as a fixed effect led to a better model fit.*

| | LME with all raters (pre-registered) | | LME without Rater 4 (exploratory) | | LME with only poor-quality scans (exploratory) | |
|---|---|---|---|---|---|---|
| Model | baseline | alternative | baseline | alternative | baseline | alternative |
| sample size | 1480 | 1480 | 1110 | 1110 | 152 | 152 |
| params | 4 | 5 | 4 | 5 | 4 | 5 |
| AIC | 1546.788 | 1542.504 | 1326.116 | 1321.000 | 166.5509 | 164.1495 |
| BIC | 1567.987 | 1569.003 | 1346.164 | 1346.060 | 178.6464 | 179.2689 |
| Log-likelihood | -769.394 | -766.252 | -659.058 | -655.499 | -79.27544 | -77.07475 |
| deviance | 1538.788 | 1532.504 | 1318.116 | 1311.000 | 158.5509 | 154.1495 |

AIC: Akaike information criterion
BIC: Bayesian information criterion
params: Number of Parameters
sample size: Number of ratings considered in the model

***S7 Table. Results of likelihood ratio test comparing LMEs with and without defacing as a fixed effect.*** $p_{FDR}$ *corresponds to p-values corrected for multiple comparisons controlling for false discovery rate (Benjamini and Hochberg 1995). The correction accounted for the p-values from the three likelihood ratio tests and the three p-values related to the defaced effect in the rm-ANOVA (c.f., Table S4). The results showed that including defacing as a fixed effect explained significantly more variance in the full-dataset scenario and the one excluding Rater 4, indicating that defacing significantly influenced the manual ratings. On the contrary, the likelihood ratio test with only low ratings was not significant, which is most likely explained by the low sample size (152 ratings only; see Table S6).*

| | LME with all raters (pre-registered) | LME without Rater 4 (exploratory) | LME with only poor-quality scans (exploratory) |
|---|---|---|---|
| $\chi^2$ | 6.283544 | 7.115882 | 4.401385 |
| df | 1 | 1 | 1 |
| $\mathbb{P}(>\chi^2)$ | 0.0122 | 0.0076 | 0.036 |
| $p_{FDR}$ | 0.0183 | 0.016 | 0.0432 |
| p<.02 | * | * | |

df: degrees of freedom

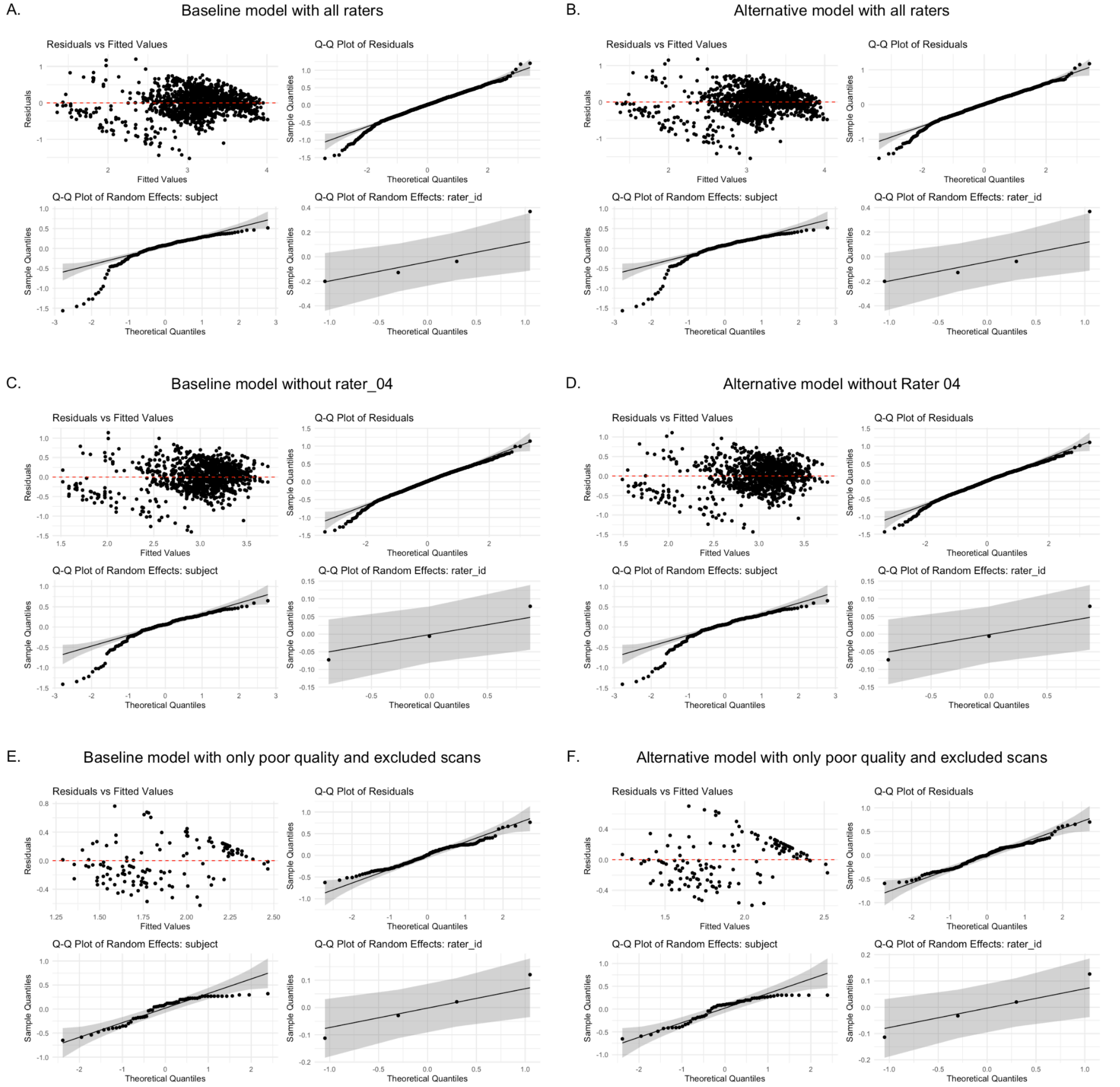

***S10 Figure. Diagnostic plots for all LMEs.*** *We report the scatter plot residuals versus fitted values and Q-Q plots (Wilk and Gnanadesikan 1968) for the residuals and the random effects included in the model to better understand model fits. Most points in the residuals versus fitted values scatter plot were evenly scattered around zero, suggesting a good fit. However, models could better fit low ratings because they showed an increasing spread as the value increased to about 2.5 for models A, B, C, and D and up to about 1.75 for models E and F. Although both latter models showed more heteroscedasticity in the residuals versus fitted values scatter plot than the other models, the overlap of all the points with the shaded gray area in the Q-Q plot indicated that the residuals were normally distributed, which is a sign of good model fit. The residuals of models A, B, C, and D deviated slightly from normality in the low ratings regime. We also present the Q-Q plots of the random effects because the latter are modeled as Gaussians in an LME; as such, the Q-Q plot allows us*

*to verify whether normality is a reasonable assumption. Modeling the rater factor as Gaussian is a good fit for all models; however, this assumption is not optimal for the subject factor in models A, B, C, and D as its distribution deviates from normality.*

***S8 Table. Variance of random effects coefficients and residuals.*** *For each LME, the residual variance was lower in the alternative model (i.e., the model including defacing as a fixed effect) than in the baseline model. Moreover, the rater variance was much lower in the LME without Rater 4 because we saw in Fig. 3 that Rater 4 presents a rating distribution considerably different from the other raters.*

| | Model | Subject (intercept) | Rater (intercept) | Residual |
|---|---|---|---|---|
| **LME with all raters** | baseline | 0.147290 | 0.049580 | 0.121690 |
| | alternative | 0.147360 | 0.049580 | 0.121100 |
| **LME without Rater 4** | baseline | 0.156950 | 0.004675 | 0.135069 |
| | alternative | 0.157123 | 0.004678 | 0.134032 |
| **LME with only poor-quality scan** | baseline | 0.105630 | 0.012680 | 0.106620 |
| | alternative | 0.112590 | 0.013400 | 0.099480 |

#### S3.3. Post-hoc power analysis of the likelihood ratio test

**Pre-registered sensitivity analysis.** The likelihood ratio test compares the ratio of likelihoods to a $\chi_2$-distribution with a degree of freedom (df) equal to the difference in parameter counts of the nested full and reduced models (df=1 in our case) (Gudicha, Schmittmann, and Vermunt 2016; Li, Zhang, and Dai 2009). As such, a proxy for effect size is given by the noncentrality parameter (Li, Zhang, and Dai 2009). The noncentrality parameter associated with the likelihood ratio test was 13.017 (Fig. S12), and a larger value can be interpreted as a larger power. To the best of our knowledge, no reference scale exists to interpret the noncentrality parameter's value, in contrast to the one that exists for Cohen's *d* effect size (Cohen 1988).

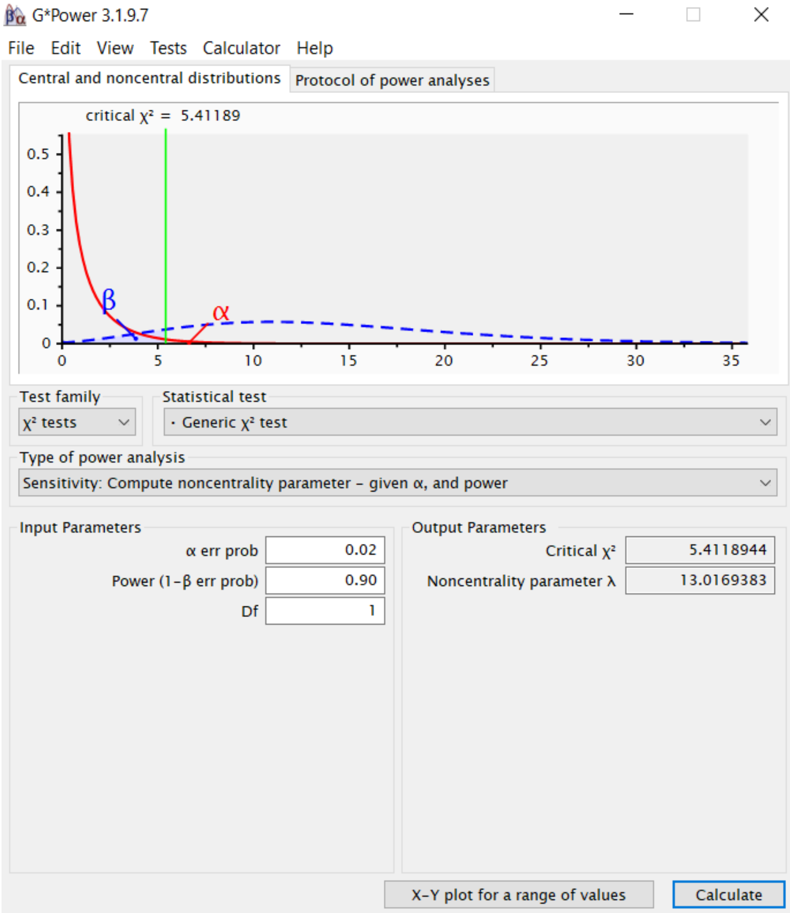

***S11 Figure. The noncentrality parameter associated with the likelihood ratio test, a proxy for its power (Li, Zhang, and Dai 2009), is 13.017.*** *A larger value is equivalent to a larger power. We ran this sensitivity analysis with G*Power (Faul et al. 2009).*

**Post-hoc power analysis.** The noncentrality parameter related to the likelihood ratio tests once the linear mixed-effect models were fitted to the data were computed using the unbiased estimation proposed in Li, Zhang, and Dai (2009):

$$\hat{\lambda}_{\beta}(x) \; = \; \max\{X - df, \beta X\},\; 0 < \beta \le 1, \quad \text{(Equation S6)}$$

where $X$ corresponds to the chi-square statistic of the likelihood ratio test and assuming $\beta = \frac{1}{1+df}$. The results of the post-hoc power analysis are presented in Table S9 below.

***S9 Table. Noncentrality parameter.*** *Noncentrality parameter estimated for each LME using Equation S6.*

| | LME with all raters (pre-registered) | LME without Rater 4 (exploratory) | LME with only poor-quality scans (exploratory) |
|---|---|---|---|
| χ2 | 6.28 | 7.11 | 4.40 |
| df | 1 | 1 | 1 |
| $\hat{\lambda}_{1/1+df}(\chi 2)$ | 5.28 | 6.11 | 3.40 |

### S4. Analysis of the Image Quality Metrics (IQMs)

#### S4.1. IQMs distribution

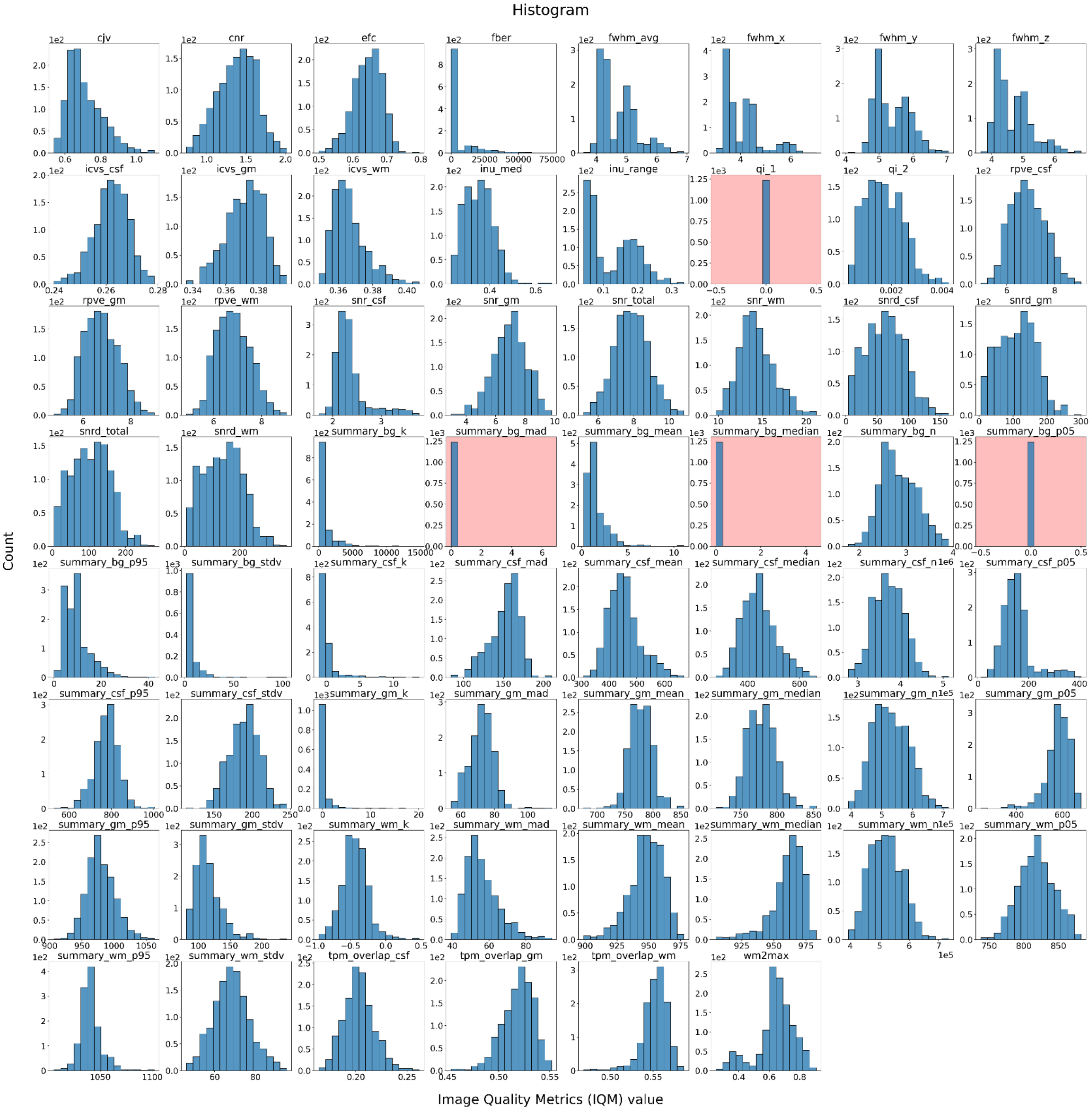

***S12 Figure. Distribution of the IQMs.*** *The IQMs calculated from defaced and nondefaced images were pooled together to plot the distributions. These plots show that the Mortamet quality index 1 (qi_1; Mortamet et al. 2009) and the 5$^{th}$ percentile of the background intensity (summary_bg_p05) were problematically constant at 0. The median of the background intensity (summary_bg_median) and the median absolute deviation of the background intensity (summary_bg_mad) were always zero except for two images, which correspond to the defaced and nondefaced images of the same subject. These observations led us to exclude those IQMs from our analysis.*

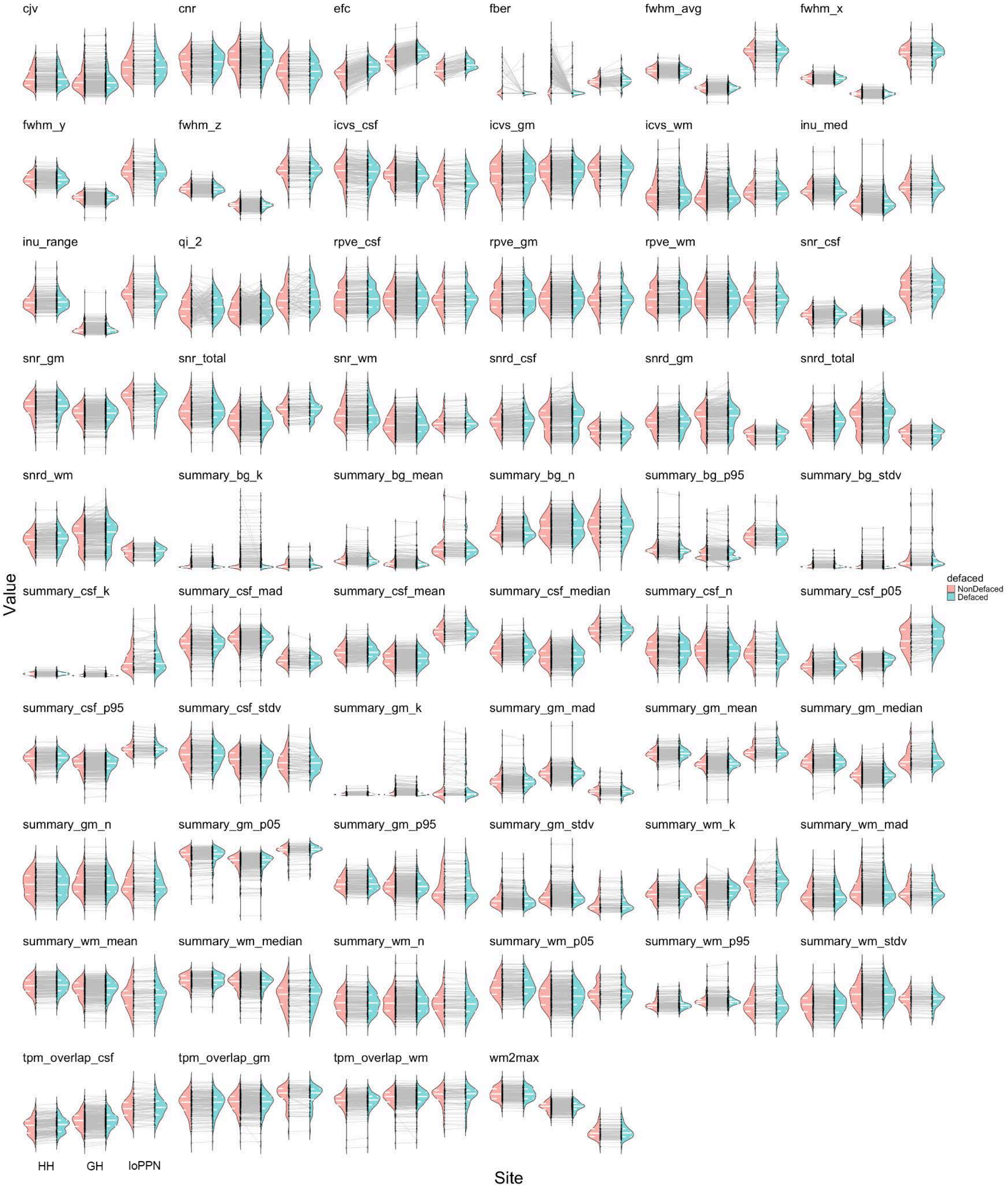

***S13 Figure. IQMs distribution per site and condition.*** *The gray lines highlight the evolution of the IQM computed from the nondefaced image to its defaced counterpart. The full white line in the violintplot represents the distribution median, while the dashed white lines represent the 25% and 75% quantiles. Many IQMs such as the full width at half maximum (fwhm) variants, the white-matter to maximum intensity ratio (wm2max), or the range of the intensity non-uniformity*

(inu_range), were distributed differently across various sites, indicating a clear site-effect. This prompted our attention to better mitigating the site-effect in the standardization and PCA. The evolution lines indicated that most IQMs were stable before and after defacing except for entropy-focus criterion (efc) and foreground-to-background energy ratio (fber) that showed clear defacing bias. Mortamet's quality index 2 (qi_2; Mortamet et al. 2009) also showed that the value of the IQM varied after defacing, but the variation was not consistent across participants; that is, some participants' quality index increased, and others decreased, such that the overall distribution remained similar.*

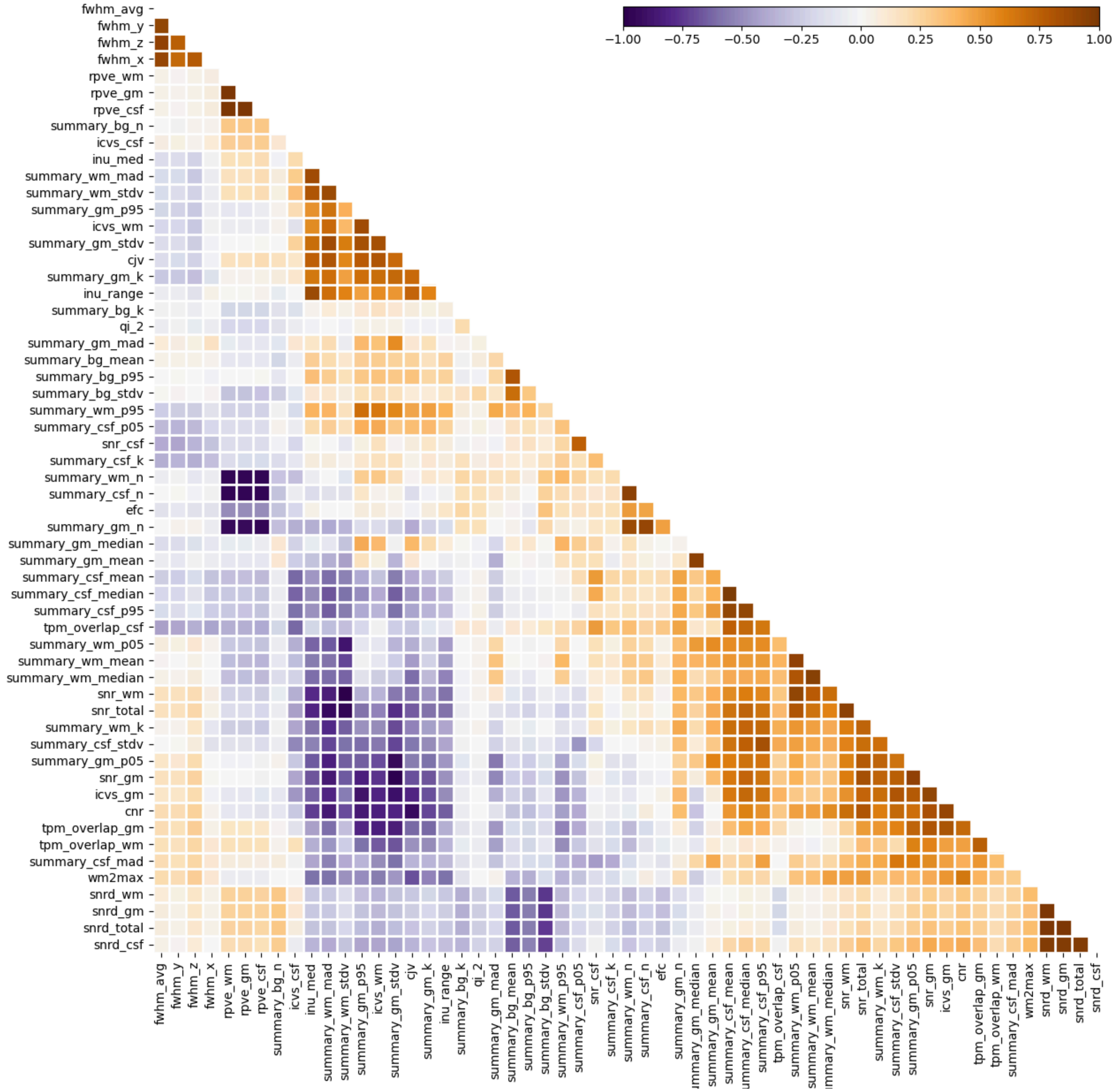

***S14 Figure. Several IQMS were highly correlated**. This plot shows the correlation between the 58 IQMs used in this study. We performed hierarchical clustering on the correlation plot to visualize the clusters of correlated IQMs more clearly.*

#### S4.2. Bland-Altman plots

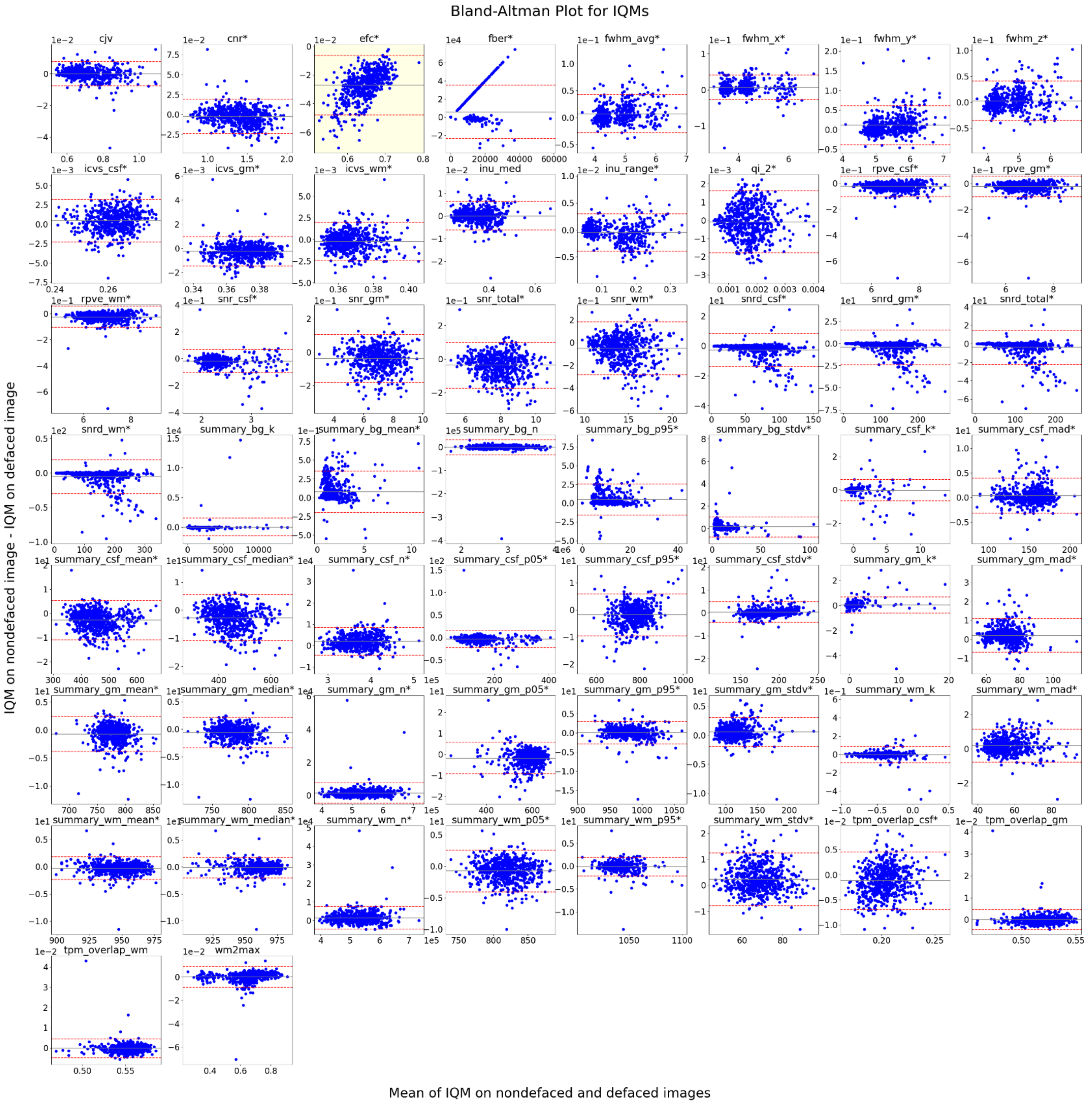

***S15 Figure. Bland-Altman plots for all non-excluded IQMs.*** *Only the entropy-focus criterion (efc) IQM presented a bias that was consistently present in all samples and consistently in the same direction as indicated by the 95% limits of agreement that did not include zero (highlighted in yellow). The bias, shown as the gray line, was computed as the mean of the differences. A bias of 0 indicates perfect agreement. A bias was considered significant when its 95% CI did not contain zero, and significance is marked by a star next to the IQM name. The 95% CI was calculated as bias ± 1.96 ** $\frac{SD}{\sqrt{n}}$ *where* $SD$ *is the standard deviation of the differences between image pairs and* $n$ *is the sample size. We computed both parametric and non-parametric 95% CIs. The parametric 95% CI was calculated as bias ± 1.96 ** $\frac{SD}{\sqrt{n}}$ *where* $SD$ *is the standard deviation of the differences between image pairs and* $n$ *is the sample size. We computed the non-parametric 95% CI using bootstrapping. A star next to the IQM name indicated significant bias, with both methods yielding*

*consistent results. Because of our large sample size ($n$=619), the 95% CI are very narrow, making most of the biases significant despite being close to zero. For legibility, the 95% CI were not shown here but were reported in a CSV file on Github[1]. The 95% limits of agreement, shown as dashed red lines, were calculated as bias ± 1.96* $SD$. These limits indicate the range where 95% of differences between the two conditions are expected to fall, reflecting their agreement. The bias 95% CI quantifies the precision of the estimated bias, showing the range in which the actual bias is likely to lie based on the sample.*

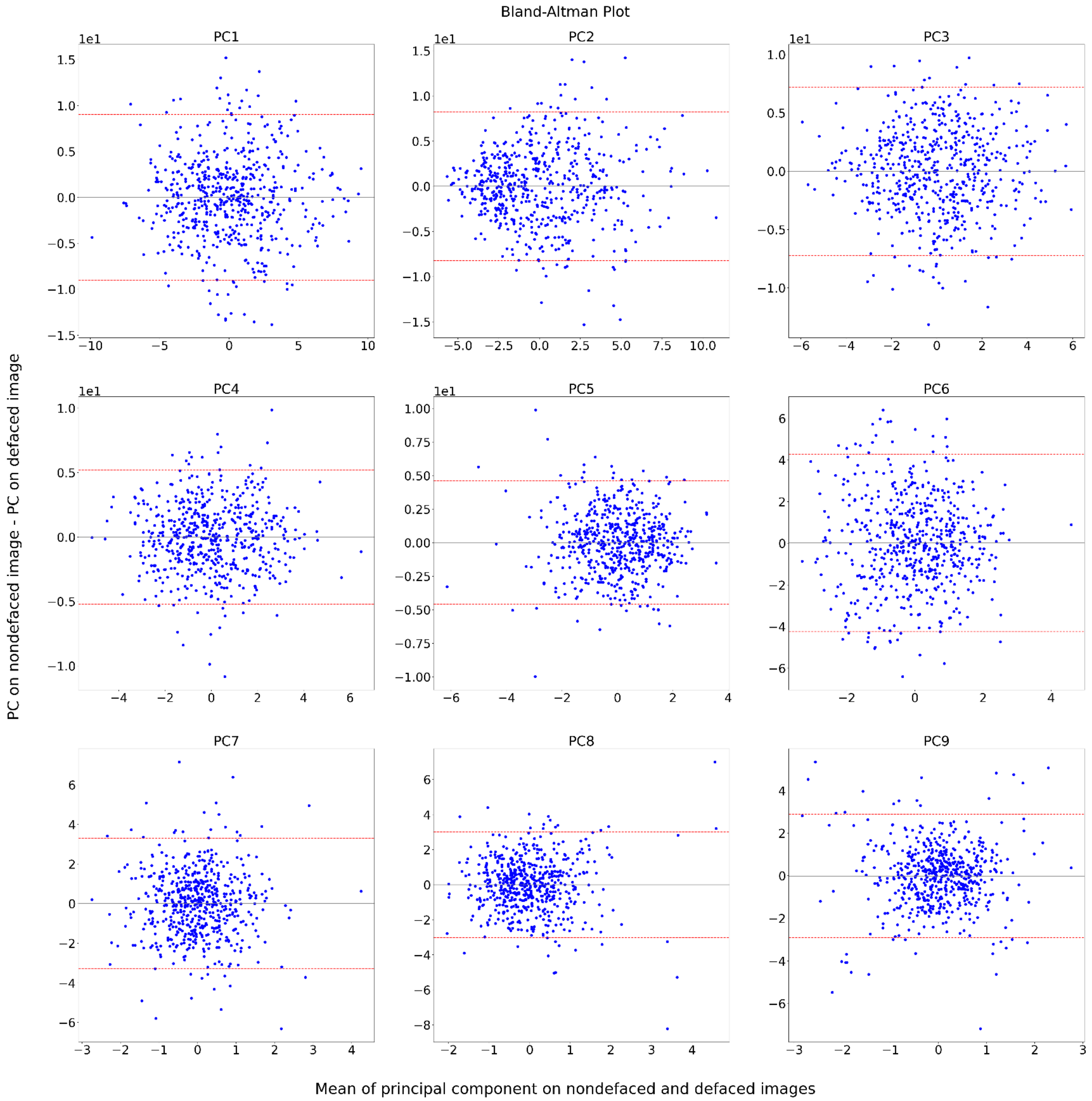

***S16 Figure. Bland-Altman plots for all principal components extracted with a single standardization and PCA step.*** *No principal component presented a significant bias between the defaced and nondefaced images because the 95% CI (computed and reported as indicated in Fig. S16) contained 0 for all principal components. The BA plots are constructed the same way as in Fig. S16.*

[1] https://github.com/TheAxonLab/defacing-and-qc-analysis/tree/main/statistical_analysis/IQMs/bias_confidence_intervals.csv

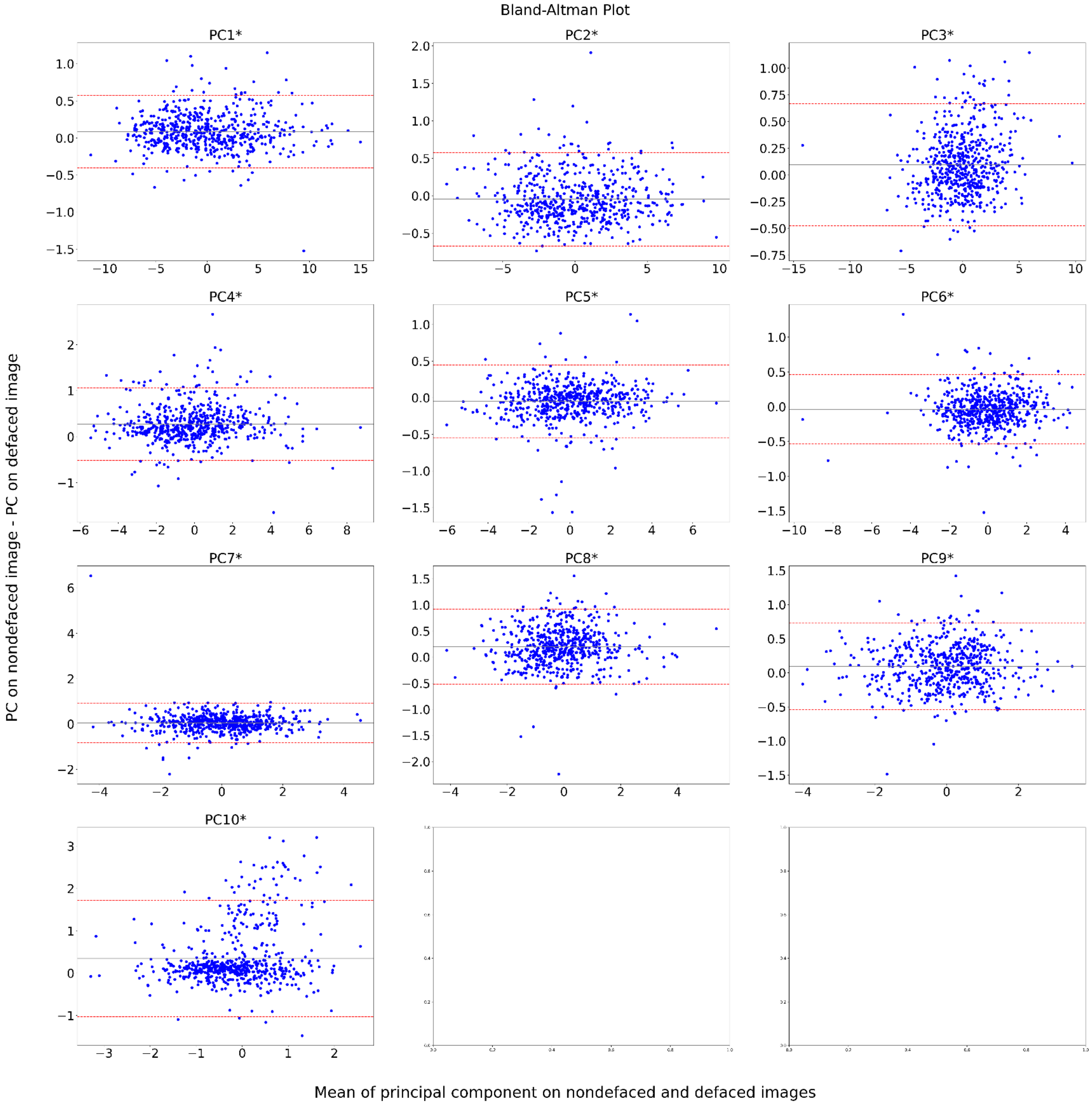

***S17 Figure. Bland-Altman plots for all principal components extracted with standardization per site but a single PCA step.*** *As explained in the caption of Fig. S16, our large sample size (n=619) results in a very narrow 95% CI, making all biases statistically significant despite being close to zero. The BA plots are constructed the same way as in Fig. S16.*

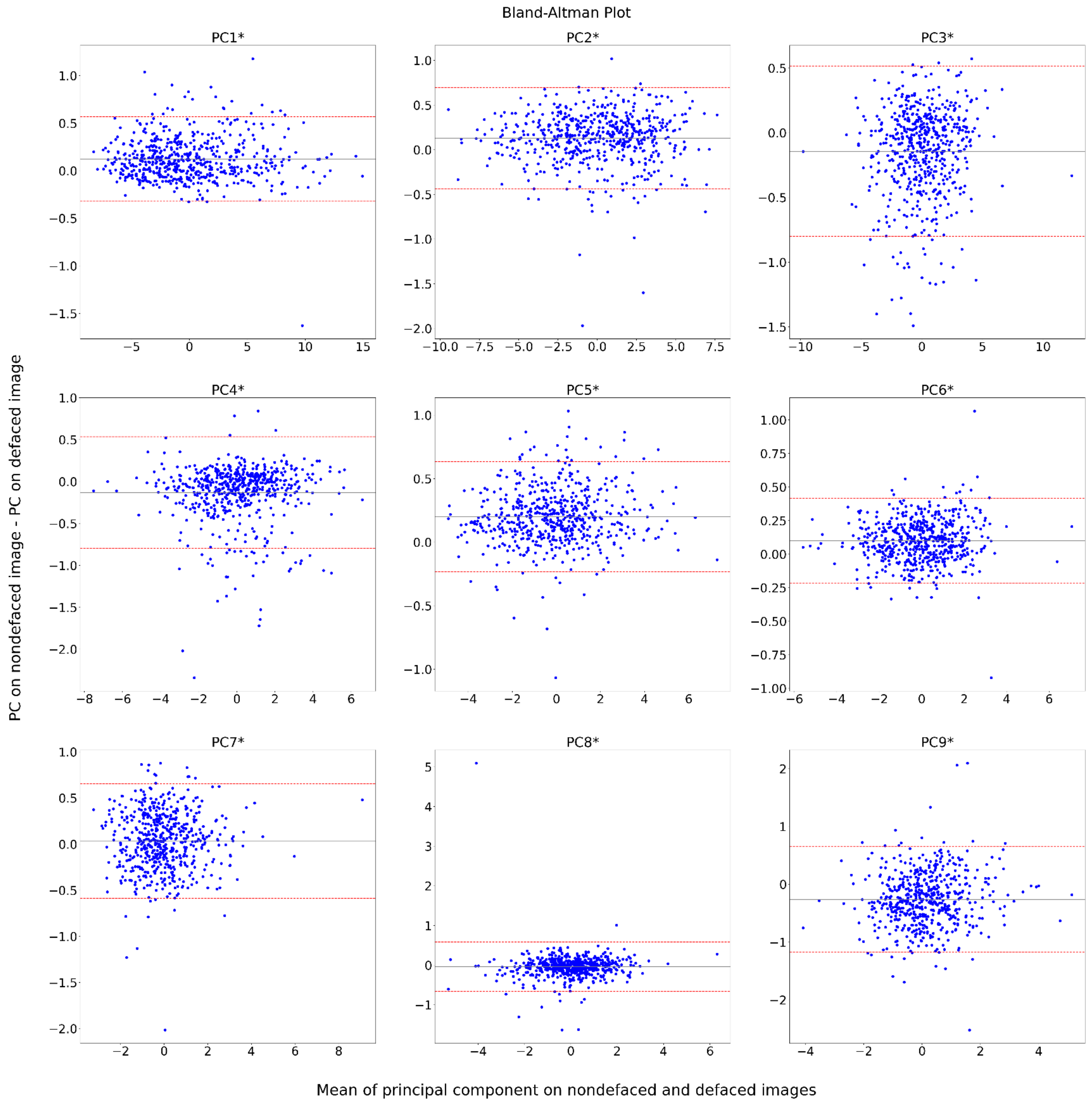

***S18 Figure. Bland-Altman plots for all principal components extracted with standardization and PCA per site.*** *Like Fig. S18, all biases were significant despite being close to zero because of our large sample size ($n$=619). The BA plots are constructed the same way as in Fig. S16.*

#### S4.3. Statistical analyses

##### S4.3.1. Dimensionality reduction

***S10 Table. Eigenvalues associated with the principal components.*** *Following the Kaiser criterion, we kept the first nine components in the scenario with a single standardization and a single PCA, and when the IQMs were standardized per site and a single PCA. We retained the first ten components for the scenario with standardization and PCA per site.*

| | Standardization and PCA per site | | | Standardization per site and single PCA | Single standardization and PCA |
|---|---|---|---|---|---|
| | Site 0 | Site 1 | Site 2 | | |
| PC1 | 19.28 | 18.95 | 14.95 | 17.70 | 14.75 |
| PC2 | 10.63 | 9.32 | 12.38 | 9.83 | 13.10 |
| PC3 | 5.11 | 5.94 | 6.72 | 5.22 | 7.64 |
| PC4 | 4.51 | 4.17 | 4.94 | 4.08 | 4.87 |
| PC5 | 4.09 | 3.68 | 4.24 | 3.99 | 3.12 |
| PC6 | 2.25 | 2.45 | 3.01 | 2.62 | 2.74 |
| PC7 | 1.85 | 1.57 | 2.25 | 1.81 | 1.43 |
| PC8 | 1.43 | 1.54 | 1.66 | 1.71 | 1.31 |
| PC9 | 1.36 | 1.33 | 1.33 | 1.37 | 1.17 |
| PC10 | 1.09 | 1 | 1.14 | 0.95 | 0.88 |
| PC11 | 0.91 | 0.91 | 0.78 | 0.92 | 0.85 |
| PC12 | 0.76 | 0.88 | 0.75 | 0.81 | 0.80 |
| PC13 | 0.7 | 0.83 | 0.65 | 0.76 | 0.75 |
| PC14 | 0.56 | 0.76 | 0.56 | 0.75 | 0.70 |
| PC15 | 0.47 | 0.66 | 0.44 | 0.67 | 0.62 |
| PC16 | 0.44 | 0.57 | 0.37 | 0.56 | 0.52 |
| PC17 | 0.4 | 0.46 | 0.31 | 0.52 | 0.45 |
| PC18 | 0.34 | 0.41 | 0.26 | 0.42 | 0.39 |
| PC19 | 0.3 | 0.39 | 0.23 | 0.40 | 0.35 |
| PC20 | 0.24 | 0.36 | 0.17 | 0.39 | 0.28 |
| PC21 | 0.22 | 0.3 | 0.15 | 0.36 | 0.23 |
| PC22 | 0.2 | 0.25 | 0.11 | 0.30 | 0.22 |
| PC23 | 0.17 | 0.24 | 0.1 | 0.27 | 0.16 |
| PC24 | 0.13 | 0.19 | 0.1 | 0.23 | 0.14 |
| PC25 | 0.11 | 0.18 | 0.09 | 0.20 | 0.11 |
| PC26 | 0.1 | 0.14 | 0.07 | 0.16 | 0.08 |
| PC27 | 0.07 | 0.13 | 0.06 | 0.14 | 0.06 |
| PC28 | 0.06 | 0.08 | 0.03 | 0.11 | 0.05 |
| PC29 | 0.05 | 0.06 | 0.03 | 0.09 | 0.04 |

| | Standardization and PCA per site | | | Standardization per site and single PCA | Single standardization and PCA |
|---|---|---|---|---|---|
| | Site 0 | Site 1 | Site 2 | | |
| PC30 | 0.03 | 0.06 | 0.02 | 0.07 | 0.03 |
| PC31 | 0.03 | 0.05 | 0.02 | 0.06 | 0.03 |
| PC32 | 0.02 | 0.04 | 0.02 | 0.05 | 0.02 |
| PC33 | 0.02 | 0.03 | 0.01 | 0.04 | 0.02 |
| PC34 | 0.01 | 0.02 | $9.57 \cdot 10^{-3}$ | 0.04 | 0.02 |
| PC35 | 0.01 | 0.01 | $7.11 \cdot 10^{-3}$ | 0.03 | 0.01 |
| PC36 | 0.01 | 0.01 | $6.52 \cdot 10^{-3}$ | 0.03 | 0.01 |
| PC37 | $7. \cdot 10^{-3}$ | $7 \cdot 10^{-3}$ | $5.50 \cdot 10^{-3}$ | 0.02 | 0.01 |
| PC38 | $6. \cdot 10^{-3}$ | $6.55 \cdot 10^{-3}$ | $4.11 \cdot 10^{-3}$ | 0.02 | 0.01 |
| PC39 | $4. \cdot 10^{-3}$ | $5.65 \cdot 10^{-3}$ | $3.25 \cdot 10^{-3}$ | 0.02 | 0.01 |
| PC40 | $3. \cdot 10^{-3}$ | $4.76 \cdot 10^{-3}$ | $2.28 \cdot 10^{-3}$ | 0.01 | 0.01 |
| PC41 | $20. \cdot 10^{-3}$ | $3.96 \cdot 10^{-3}$ | $1.52 \cdot 10^{-3}$ | 0.01 | 0.01 |
| PC42 | $1 \cdot 10^{-3}$ | $3.25 \cdot 10^{-3}$ | $8.79 \cdot 10^{-4}$ | 0.01 | $4 \cdot 10^{-3}$ |
| PC43 | $1.17 \cdot 10^{-3}$ | $1.65 \cdot 10^{-3}$ | $7.64 \cdot 10^{-4}$ | 0.01 | $3 \cdot 10^{-3}$ |
| PC44 | $7.77 \cdot 10^{-4}$ | $1.03 \cdot 10^{-3}$ | $4.89 \cdot 10^{-4}$ | 0.01 | $2 \cdot 10^{-3}$ |
| PC45 | $5.90 \cdot 10^{-4}$ | $7.27 \cdot 10^{-4}$ | $3.01 \cdot 10^{-4}$ | $4. \cdot 10^{-3}$ | $8 \cdot 10^{-4}$ |
| PC46 | $5.46 \cdot 10^{-4}$ | $5.79 \cdot 10^{-4}$ | $1.46 \cdot 10^{-4}$ | $3. \cdot 10^{-3}$ | $7 \cdot 10^{-4}$ |
| PC47 | $3.94 \cdot 10^{-4}$ | $4.66 \cdot 10^{-4}$ | $1.17 \cdot 10^{-4}$ | $3. \cdot 10^{-3}$ | $6 \cdot 10^{-4}$ |
| PC48 | $2.74 \cdot 10^{-4}$ | $3.38 \cdot 10^{-4}$ | $9.40 \cdot 10^{-5}$ | $2. \cdot 10^{-3}$ | $5 \cdot 10^{-4}$ |
| PC49 | $2.35 \cdot 10^{-4}$ | $2.90 \cdot 10^{-4}$ | $6.02 \cdot 10^{-5}$ | $2. \cdot 10^{-3}$ | $4 \cdot 10^{-4}$ |
| PC50 | $1.34 \cdot 10^{-4}$ | $2.68 \cdot 10^{-4}$ | $4.23 \cdot 10^{-5}$ | $1. \cdot 10^{-3}$ | $3 \cdot 10^{-4}$ |
| PC51 | $6.39 \cdot 10^{-5}$ | $8.74 \cdot 10^{-5}$ | $3.06 \cdot 10^{-5}$ | $9. \cdot 10^{-4}$ | $1 \cdot 10^{-4}$ |
| PC52 | $4.70 \cdot 10^{-5}$ | $4.70 \cdot 10^{-5}$ | $1.82 \cdot 10^{-5}$ | $6. \cdot 10^{-4}$ | $9 \cdot 10^{-5}$ |
| PC53 | $4.46 \cdot 10^{-7}$ | $5.37 \cdot 10^{-7}$ | $5.76 \cdot 10^{-7}$ | $4. \cdot 10^{-4}$ | $2 \cdot 10^{-6}$ |
| PC54 | $3.62 \cdot 10^{-7}$ | $3.31 \cdot 10^{-7}$ | $1.95 \cdot 10^{-7}$ | $2. \cdot 10^{-4}$ | $1 \cdot 10^{-6}$ |
| PC55 | $2.64 \cdot 10^{-29}$ | $2.47 \cdot 10^{-29}$ | $8.39 \cdot 10^{-30}$ | $1. \cdot 10^{-4}$ | $3 \cdot 10^{-29}$ |
| PC56 | $4.69 \cdot 10^{-30}$ | $7.64 \cdot 10^{-30}$ | $2.07 \cdot 10^{-30}$ | $5. \cdot 10^{-6}$ | $1 \cdot 10^{-30}$ |
| PC57 | $1.25 \cdot 10^{-30}$ | $1.24 \cdot 10^{-30}$ | $1.95 \cdot 10^{-30}$ | $2. \cdot 10^{-6}$ | $7 \cdot 10^{-31}$ |
| PC58 | $3.92 \cdot 10^{-31}$ | $3.49 \cdot 10^{-31}$ | $2.67 \cdot 10^{-31}$ | $1. \cdot 10^{-6}$ | $4 \cdot 10^{-31}$ |

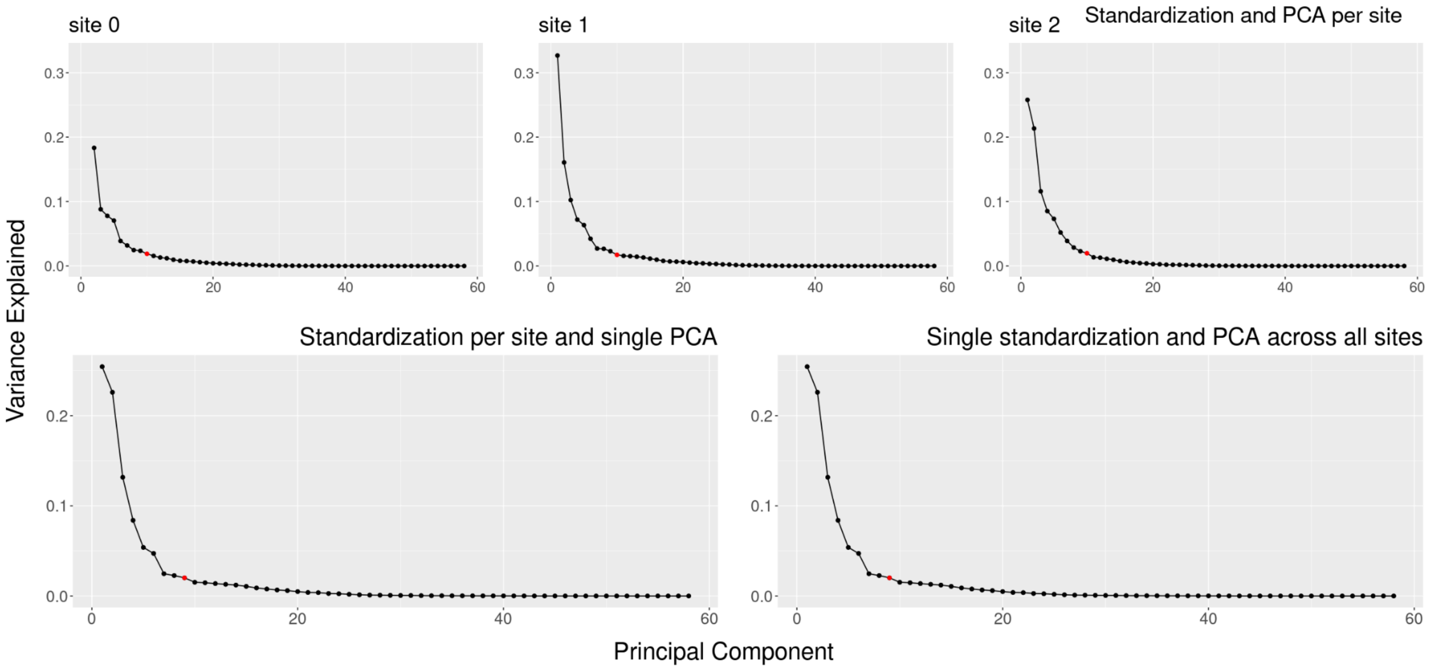

***S19 Figure. Scree plot: Variance explained by the principal components.*** *The point in red indicates the number of components the Kaiser criterion selects. The number of components chosen according to the Kaiser criterion is reasonable for all PCA as choosing one component more does not explain substantially more variance for any of the scree plots.*

##### S4.3.2. rm-MANOVA after dimensionality reduction via PCA

***S11 Table. Results of repeated-measures MANOVA on the projected IQMs.*** *Effect size is reported as partial eta-squared ($\eta_p^2$) and was computed with the function F_to_eta2 from the R package effectsize (Ben-Shachar, Lüdecke, and Makowski 2020), which implements the formula given in Equation S4 with df = numDF and df_error = denDF. $p_{corrected}$ correspond to p-values controlled for false discovery rate (Benjamini and Hochberg 1995). The pre-registered plan found neither an effect of the site nor an effect of defacing on the principal components extracted from the IQMs. However, applying the IQMs standardization and PCA separately per site mitigated the site-effects, revealing the defacing bias. The rm-MANOVA with only a small subset of IQMs (that is efc, cnr, cjv, fber and $snr_{wm}$) also revealed a significant influence of defacing on those selected IQMs.*

| | | **Wald-type Statistic (WTS)** | | | | **Modified ANOVA-type Statistic (MATS)** | | **Resampling version of the tests** | |
|---|---|---|---|---|---|---|---|---|---|
| | | Test Statistic | df | p | $p_{corrected}$ | Test Statistic | $\eta_p^2$ | paramBS (WTS) | paramBS (MATS) |
| Standardization and PCA per site (exploratory) | site | 0.358 | 2 | 0.836 | 0.994 | 0.710 | 0.001 | 0.833 | 0.833 |
| | defaced | 181.182 | 1 | <.001 | <.003 ** | 2.726 | 0.005 | <.001 *** | <.001 *** |
| Standardization per site and single PCA (exploratory) | site | 0.011 | 2 | 0.994 | 0.994 | 0.023 | $4 \cdot 10^{-5}$ | 0.995 | 0.995 |
| | defaced | 1.038 | 1 | 0.308 | 0.462 | 0.005 | $9 \cdot 10^{-6}$ | 0.31 | 0.311 |
| Single standardization and PCA across all sites (pre-registered) | site | 0.376 | 2 | 0.828 | 0.994 | 0.457 | $8 \cdot 10^{-4}$ | 0.829 | 0.827 |
| | defaced | 0.003 | 1 | 0.96 | 0.96 | 0.002 | $3 \cdot 10^{-6}$ | 0.961 | 0.961 |
| Selected subset of IQMs (exploratory) | site | 332.984 | 2 | <.001 *** | | 473.708 | 0.45 | <.001 *** | <.001 *** |
| | defaced | 59.053 | 1 | <.001 *** | | 42.246 | 0.068 | <.001 *** | <.001 *** |

#### S4.4. Defacing did not influence the IQMs' reliability (unplanned exploratory analysis)

As part of our broader investigation, we evaluated the effect of defacing on the IQMs' reliability. For every IQM, we calculated two coefficients of variation (CoV): one considering IQMs from nondefaced images, and the other from defaced images. We then tested whether the CoV distributions from both defacing conditions are significantly different from each other using a paired t-test. We also repeated the test using only the subset of IQMs selected in the rm-MANOVA analysis, namely entropy-focus criterion (efc), contrast-to-noise ratio (cnr), coefficient of joint variation (cjv), fber, and signal-to-noise ratio within white matter mask ($snr_{wm}$) (see section S3.3). Unsurprisingly, after assessing that the IQMs extracted by *MRIQC* are insensitive to defacing by design, our results support that defacing did not undermine the reliability of IQMs (see Fig. S21 for detailed results and interpretation).

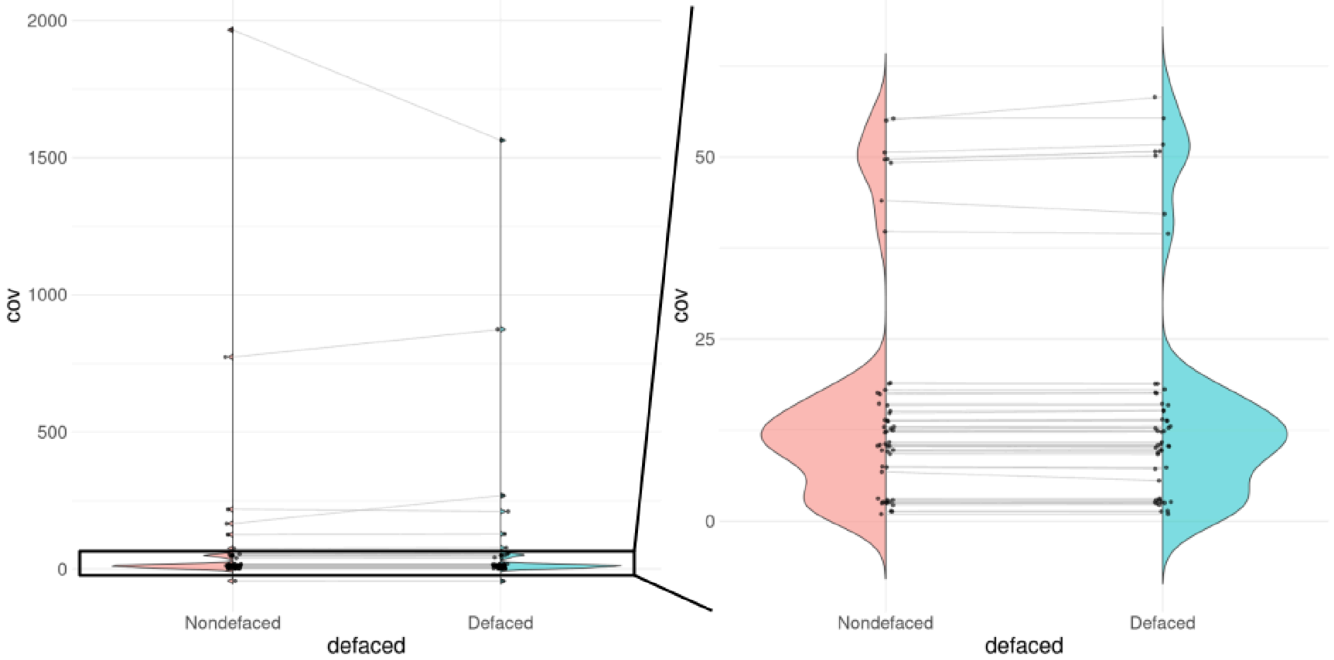

***S20 Figure. Defacing did not influence the IQMs' reliability.*** *For every IQM, we computed two coefficients of variation (CoV): one considering IQMs from nondefaced images and the other IQMs from defaced images. We then compared the distributions of these CoVs using a combination of a violin plot and a line plot. The distribution appeared very similar, and the evolution lines were majoritarily flat, showing that the IQMs reliability did not change following the defacing process. A paired t-test comparing the CoV distributions from both defacing conditions confirmed that they were not significantly different, t(57) = 0.92, p = 0.35, d=0.06. Considering only the subset of IQMs selected in the rm-MANOVA analysis (that is, efc, cnr, cjv, fber, and* $snr_{wm}$*) also revealed no significant differences in the CoV distributions, t(4) = 0.98, p=0.38, d=0.4. These results aligned with our understanding that MRIQC, by design, does not account for areas typically affected by defacing in the computation of the IQMs.*

#### S5. Importance of the background in QA/QC

During the human rater training session, we highlighted the importance of considering the background when assessing image quality. We demonstrated this by showing several examples where artifacts impacting signal of interest were more noticeable in the background, including one example shown in Fig. S22. Many of the exclusion criteria we asked the raters to consider used the background to help detect artifactual signals, as outlined in our QC protocol (Provins et al. 2023). Artifacts often stand out more in the background due to the absence of a signal of interest. While they may also affect regions of interest, their presence is more challenging to perceive in those busy areas.

***S21 Figure. Screening the background helps detect artifacts within the brain.*** *This figure (next full page) illustrates how including information outside the brain outline improves artifact detection, particularly for wrap-around artifacts. In this example, the posterior of the head is "wrapped around" and aliases over the prefrontal cortex, compromising the signal in that region. When the brain has been extracted (A), the overlap of the skull becomes elusive. Conversely, when the assessment is performed before brain extraction (B), the overlap is easily traceable, regardless of the observer's experience. All images are screenshots from MRIQC visual reports extracted from the T1w scan of subject "10377" from the Consortium for Neuropsychiatric Phenomics dataset (Gorgolewski, Durnez, and Poldrack 2017).*

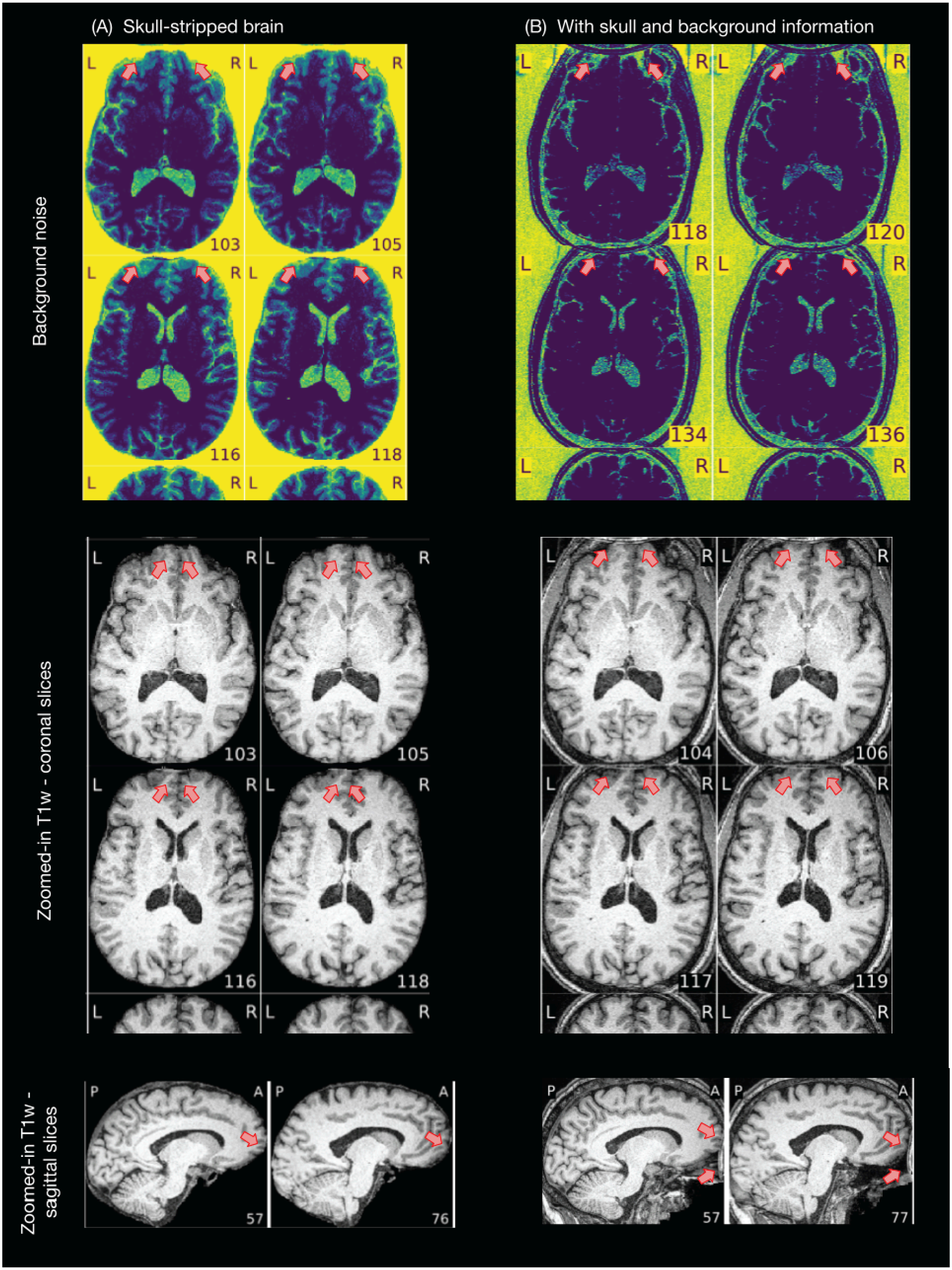
(A) Skull-stripped brain
(B) With skull and background information
Background noise
Zoomed-in T1w - coronal slices
Zoomed-in T1w - sagittal slices
L
R
P
A
103
105
116
118
118
120
134
136
103
105
116
118
104
106
117
119
57
76
57
77
